# Computational Mechanisms of Temporal Anticipation in Perception and Action

**DOI:** 10.1101/2025.03.28.646008

**Authors:** Gal Vishne, Leon Y. Deouell, Ayelet N. Landau

## Abstract

To interact effectively with our surroundings, we rely on strategies to reduce uncertainty. One important source of information is temporal regularities, which enable us to form predictions about when events will occur, and through this, prepare for them in advance. Such preparation was shown to facilitate motor planning, yet the impact of temporal anticipation on perceptual acuity is unknown, and the cognitive computations underlying this process remain debated. To answer these questions, we administered a new difficult change discrimination task with variable anticipatory periods (N=142). We show that both perceptual sensitivity and motor responses are influenced by temporal structure. Using computational modelling, we show that the cognitive operation underlying this behavior is based on a logarithmic transformation of the event hazard rate (HR) and reveal a crucial role of temporal-estimation noise in shaping this computation, both when temporal information is encoded, and when it is decoded. Together, our results highlight the fundamental contribution of anticipation in directing behavior and advance our understanding of temporal processing in the brain.

## Introduction

We live in a dynamic world, with constant sensory changes. Being able to anticipate the timing of these changes is therefore critical for effective interaction with our environment. To do so, humans (and other animals) have a remarkable ability to extract and leverage temporal structures and regularities in their surroundings.^1–6^ Temporal structure is present on a continuum: at one end, events are fully predictable, occurring precisely when expected; while at the other end, events are completely surprising, offering no opportunity for preparation. Most real-world experiences fall somewhere between these two extremes – we have some, but not all, information about when an event will occur. In these cases, we can conceive of the timing of the event as drawing from a temporal probability distribution, spanning a range of possible times. These distributions can be learned through experience and adjusted based on multiple cues. For example, when entering a bus stop, we can usually estimate that the bus will arrive within a few minutes, but often not precisely when. The assigned probability of the bus arriving at any given time is dependent on a range of factors, such as our prior experience with this bus route, or the time of day, which informs us whether we should use the distribution of buses for peak-hours or off-hours.

Experimentally, this type of anticipatory temporal information has been traditionally studied using variants of the ‘set-go’, also known as variable ‘foreperiod’ paradigms:^7–14^ First, a ‘set’ stimulus is presented (S1), then, after a variable amount of time known as the ‘foreperiod’ (FP), a ‘go’ or ‘target’ stimulus is presented (S2) and participants are required to respond to it as soon as possible. In this setup, S1 serves as a temporal reference point for participants, and indicates to them to start preparing for the ‘go’ signal. The distribution of preparatory periods (the time interval between the two stimuli), is referred to as the FP distribution. Often, there are a certain number of trials where S2 is omitted, which serve as ‘catch trials’, to ensure that participants indeed wait for the go signal before they act, rather than acting prematurely based purely on anticipation, like a runner who commits a false start. By using this type of task with variable FP distributions, it was shown that people (and other primates) are able to utilize distributional temporal information to adjust motor preparation.^7,8,10,11,13,15–20^ However, as these studies relied nearly exclusively on reaction time (RT) measures it is unclear whether perceptual acuity is also influenced by distributional temporal structures (see the <u>Discussion</u> for the relation to other forms of temporal anticipation). That is, when one can use the FP distribution to predict target times, is perceptual performance improved, and if so, what is the relationship between this improvement and the modulation of motor responses? Answering this question is the first aim of the current study.

The second aim of the current study is to shed light on the computational mechanisms which allow us to modulate behavior by environmental temporal structure, and more broadly, the mechanisms underlying temporal anticipation. To utilize temporal structure optimally, two requirements are needed: First, one needs to acquire precise information about the probability of event occurrence at each moment in time, i.e., to know the event probability density function (PDF). Second, one needs to dynamically adjust this information as time elapses and the event did not occur yet. The result of this computation is known as the hazard rate (HR) – the probability that an event will occur, given that it did not occur yet.^21^ The HR was classically assumed to underly the neural and behavioral effects of temporal anticipation, including the modulation of motor preparation by temporal probability distributions (e.g., ^2,8,21,22^). However, despite the intuitive appeal of this account, it was recently challenged on both experimental and theoretical grounds, and several alternatives have been suggested.^17–20,23^ One important issue raised by these studies is that the HR account is often taken for granted, without direct evaluation in comparison to other explanations.

To address this problem, in the current study, we construct multiple competing computational models and compare their predictions qualitatively and quantitatively (*Fig. 1*). Using this model comparison lens, we were able to examine in more detail the two underlying assumptions of the optimality account. First, it is unlikely that participants have perfect knowledge of the underlying probability distribution. One crucial factor contributing to this is the inherent noise in how brains estimate elapsed time.^24,25^ We examine the impact of temporal-estimation noise both on the formation of temporal representations (‘encoding noise’) and on how they are used to guide behavior (‘decoding noise’). We further explore the factors modulating temporal-estimation noise, testing whether it depends primarily on elapsed time (‘time-based’ modulation) or on the probability of event occurrence (‘probability-based’ modulation). Second, the HR computation itself is complex and numerically unstable, which may pose difficulties for rapid and efficient processing. Therefore, despite being the optimal computation, it may be that the brain uses approximations of this quantity for real-world behavior, for example, using the PDF itself, instead of scaling it dynamically with elapsed time.^18–20^ To test this hypothesis, we consider both the HR and the PDF as the core computation underlying event anticipation and examine several nonlinear transformations of each of these basic computations.

**Fig. 1:**
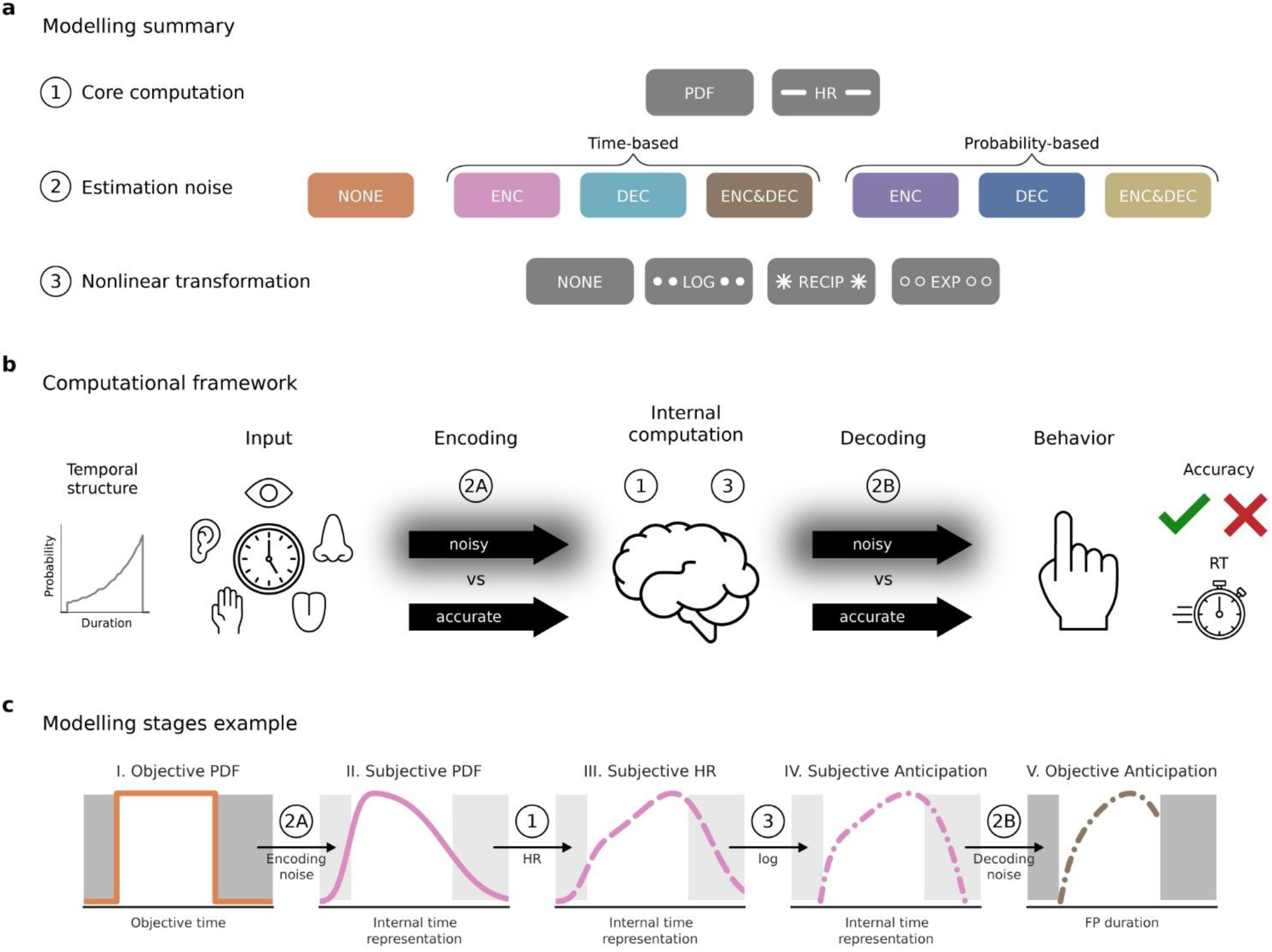
Computational modelling overview. (**a**) We examined 56 models, reflecting competing hypotheses about the algorithmic and computational steps underlying event anticipation, in three stages: (1) Core computation: event PDF (probability density function) and event HR (hazard rate; <u>Methods\Hazard rate</u>). (2) Estimation noise (<u>Methods\Temporal-estimation noise</u>): We considered separately the influence of noise on encoding and decoding of the task temporal structure (resulting in three noise models: ‘ENC’ only encoding noise, ‘DEC’ only decoding noise, ‘ENC&DEC’ encoding and decoding noise). For each noise model, we evaluated two different factors for modulating the noise, elapsed time (‘time-based’) and event probability (‘probability-based’). Together with the noise-free model (‘NONE’), this results in 7 models for each core computation. (3) Nonlinear transformation (<u>Methods\Nonlinear transformations</u>): four options for each of the models (‘NONE’: no nonlinearity, ‘LOG’: logarithm, ‘RECIP’: reciprocal, ‘EXP’: exponential). (**b**) We model the task-level transformation of external temporal structure into behavioral modulation as consisting of encoding, internal computation, and decoding. Temporal structure is captured by the task-level PDF (depicted in the left for the FLIP-EXP condition, see *Fig. 2b* for all conditions). Encoding takes us from the objective PDF to a subjective PDF (internal representation of the task structure). Anticipation is computed based on this internal representation: First, in HR-based models, the subjective PDF is transformed into a subjectively encoded HR function (modelling stage 1, see (a)), followed in some models by another nonlinear computation (modelling stage 3). Decoding is the process of employing the internal anticipation to modulate behavior (accuracy and RT, depicted in the right). The influence of temporal-estimation noise (modelling stage 2) was tested separately for encoding (2A) and decoding (2B). This panel was designed using resources from Flaticon.com. (**c**) Illustration of the influence of the modelling procedure on the UNI distribution condition. Each panel depicts one step in the computation of the logarithm of the HR model with time-based noise in encoding and decoding (HR-log-ENC&DEC-N-time), which provided the best fit for both accuracy and RT (see main text). For each intermediate stage of the computation the color and line style correspond to the model in (a) whose final output is identical to the model up to that stage. Panel I shows the objective PDF, which was used as the basis for all models. Panel II shows the influence of encoding, shown here with time-based estimation-noise. Panel III shows the result of the HR computation is shown, which increases the emphasis on long FPs. This is followed in panel IV by a nonlinear transformation (logarithm here, which accentuates the difference between high and low probability FPs). Finally, panel V shows the result of decoding this subjective anticipation, shown with time-based estimation-noise.

We developed a novel task combining visual change discrimination with variable FP paradigms (*Fig. 2a*). In contrast to standard change discrimination tasks, where both stimuli (S1, S2) are presented for the same duration, here S1 was presented for a longer time, and its presentation duration was varied on a trial-by-trial basis, drawing from one of three predefined temporal probability distributions (between participant manipulation; *Fig. 2b*). By introducing a difficult perceptual judgement to the traditional FP design, we were able to analyze the influence of temporal anticipation on both perceptual acuity and motor performance and show that both are strongly modulated by the temporal structure of the environment. Furthermore, using our detailed modelling approach, we find that the cognitive computation underlying these behavioral modulations is based described as the logarithm of the event HR. Finally, we show that this computation is further shaped by temporal-estimation noise both at the encoding stage and at the decoding stage, and that estimation noise is modulated by elapsed time, not the probability of event occurrence. Together, these findings shed light on coding of time in the brain and inform our understanding of how the brain utilizes temporal structure to guide behavior.

**Fig. 2:**
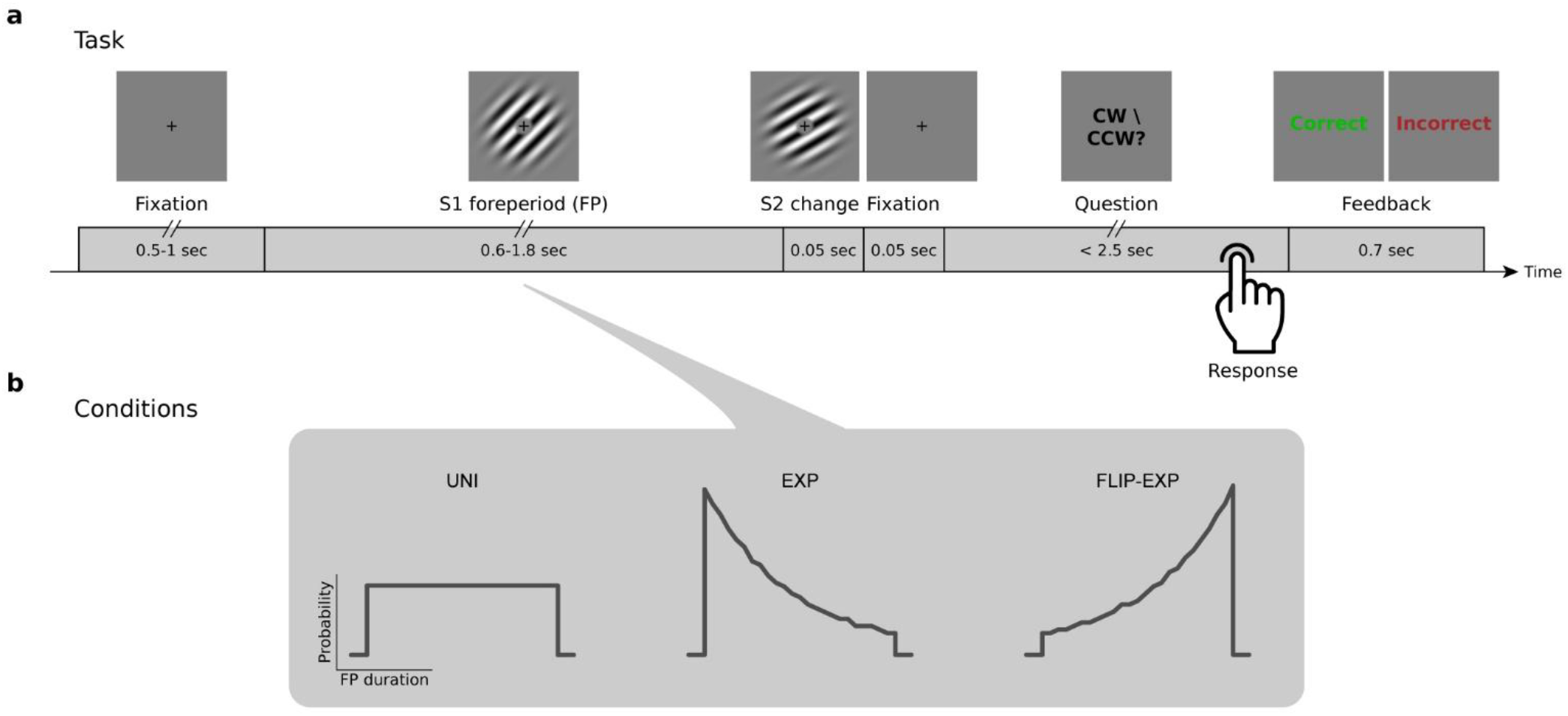
Task and conditions. (**a**) Trial design: Participants performed a change discrimination task with variable anticipatory periods. In standard trials, participants were presented with two consecutive oriented gratings (S1 and S2; see <u>Methods\Stimuli</u>), and were asked to judge the orientation change between them (clockwise or counterclockwise). S1 was presented for an extended duration, which served as the foreperiod (FP). In addition to the standard trials presented here, we included a small fraction of trials without any change, to discourage premature responses (‘no-change’ trials, ∼12.5% of all trials; see <u>Methods\Trial procedure</u> for more details). (**b**) Conditions: Participants were assigned to one of three experimental conditions, which determined the task-level distribution of S1 durations (<u>Methods\Foreperiod distributions</u>): uniform (UNI, N=47), exponential (Exp, N=41) and flipped-exponential (FLIP-EXP, N=54).

## Results

We present the results of N=142 participants (online recruitment, see <u>Methods\Participants</u> for more details about the groups and the exclusion criteria). In each trial, participants viewed two consecutive oriented gratings (S1 and S2) and were tasked with judging whether S2 was tilted clockwise or counterclockwise relative to S1 (*Fig. 2a*). The duration of S1 varied on a trial-by-trial basis, drawing in separate groups from one of three predefined temporal probability distributions (*Fig. 2b*; <u>Methods\Foreperiod distributions</u>) – uniform (UNI; N=47), exponential (EXP; N=41) and flipped-exponential (FLIP-EXP; N=54). Similar to classical FP paradigms, in a small proportion of trials S2 was omitted (‘catch trials’, <u>Methods\Trial procedure</u>). To ensure that we can identify modulation of perceptual ability in each participant, task difficulty was controlled using an ongoing staircase procedure, aimed at reaching ∼80% accuracy (<u>Methods\Staircase procedure</u>). Outlier trials with exceptionally fast or exceptionally slow RT, were removed from analysis (thresholds were defined based on individual RT distributions, see <u>Methods\Data selection</u>; results were similar without this step). Overall task performance was not significantly different between the three distribution conditions, with comparable task accuracy (mean ± SEM across participants: ACC_UNI_ 79.7±0.06%, ACC_EXP_ 79.5±0.07%, ACC_FLIP-EXP_ 79.6±0.07%; one-way ANOVA F(2,139)=1.67 p>0.19, η_p_^2^=0.02), and comparable median RT (RT_UNI_ 565.9±11 msec, RT_EXP_ 545±12.7 msec, RT_FLIP-EXP_ 549.1±11.4 msec; one-way ANOVA F(2,139)=0.86 p>0.42 η_p_^2^=0.01). Additionally, there were no significant differences in median tilt magnitude (Tilt_UNI_ 0.74±0.06°, Tilt_EXP_ 0.67±0.05°, Tilt_FLIP-EXP_ 0.68±0.05°; one-way ANOVA F(2,139)=0.5 p>0.6 η_p_^2^=0.01).

### Perceptual accuracy and reaction times are both modulated by temporal structure

To understand how the passage of time influences behavior, we examined the relation between FP duration and performance in the task, focusing first on performance in the shortest and longest FP durations. For each participant, we computed the accuracy (ACC) and the median RT (using only correct trials) separately for trials with FPs between 0.6-0.7 sec and 1.7-1.8 sec (<u>Methods\Behavioral quantification</u>). We found that accuracy was significantly higher for longer vs shorter durations in all three distributions (*Fig. 3a*; paired t-test per distribution, Bonferroni corrected for three distributions; UNI t(46)=6.68, p_Bonf_<10^−7^, Cohen’s d=0.98; EXP t(40)=3.15, p_Bonf_<0.01, d=0.5; FLIP-EXP t(53)=9.73, p_Bonf_<10^−12^, d=1.32; ACC_long_-ACC_short_ (mean ± SEM): ΔACC_UNI_ 9.2±1.4%, ΔACC_EXP_ 5.3±1.7%, ΔACC_FLIP-EXP_ 17.8±1.8%). Turning to RT (*Fig. 3b*), we found faster responses for longer vs shorter FPs in the UNI and FLIP-EXP conditions (UNI t(46)=-6.57, p_Bonf_<10^−6^, d=0.96; FLIP-EXP t(53)=-9.84, p_Bonf_<10^−12^, d=1.34; ΔRT_UNI_ −52.6±8 msec, ΔRT_FLIP-EXP_ −156.5±15.9 msec), but not in the EXP condition, where the pattern was nominally reversed, with slightly slower responses for longer durations (although this was not significant, t(40)=1.74, p_Bonf_>0.26, d=0.27; ΔRT_EXP_ 20±11.5 msec). Thus, both accuracy and reaction speed revealed a strong influence of FP duration, with overall higher accuracy and faster (or similar) RT for longer FPs.

**Fig. 3:**
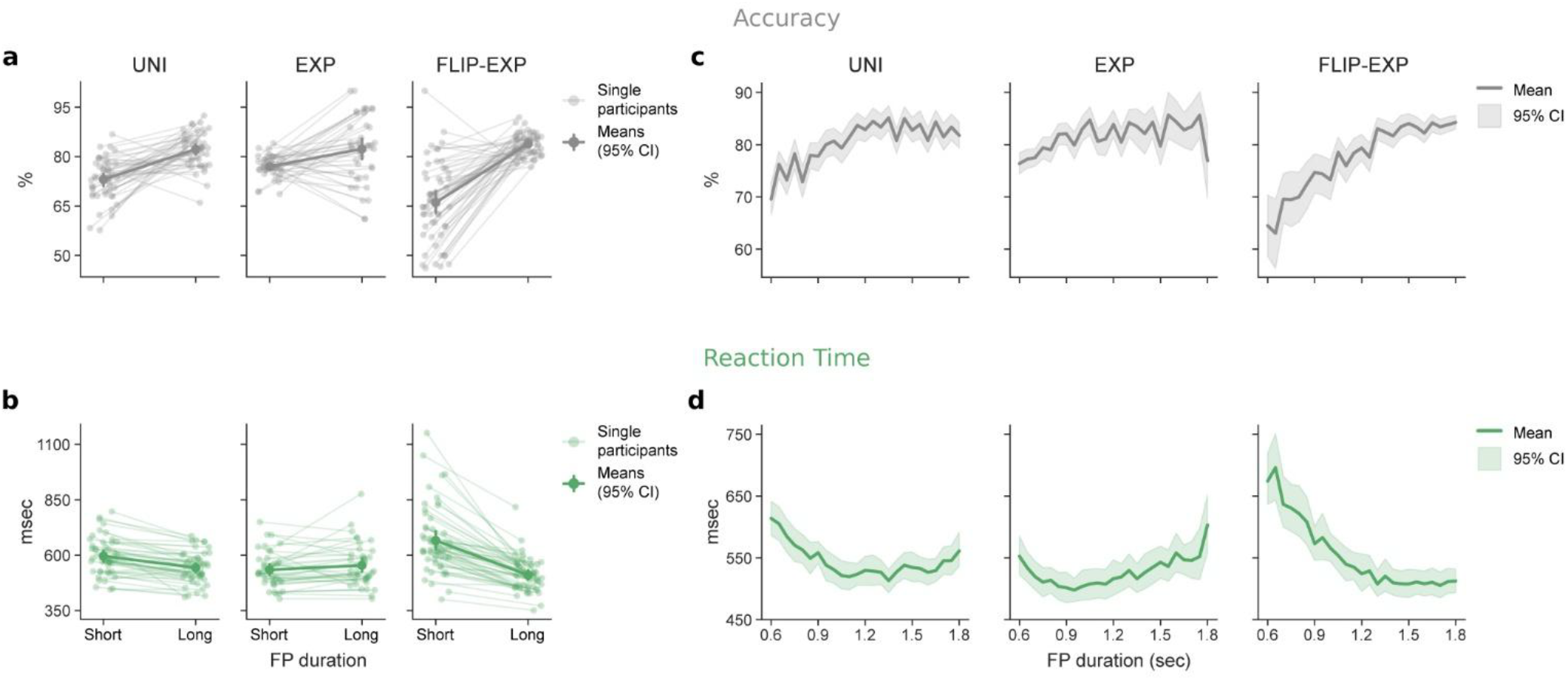
Behavioral performance in each distribution condition. (**a**-**b**) Performance in the shortest and longest FP durations (short: 0.6-0.7 sec, long: 1.7-1.8 sec). Semi-transparent dots and lines: single participant data, opaque dots and bold lines: group means. Error bars: 95% confidence interval of the mean (CI; bootstrapping, 1000 iterations). (**a**) Accuracy (percent correct trials) was significantly influenced by FP duration in all distribution conditions. (**b**) RT (median reaction time in correct trials) was significantly influenced by FP duration in the UNI and FLIP-EXP conditions but not in the EXP condition. The influence of time was strongly modulated by distribution condition for both behavioral measures (see main text). (**c**-**d**) Full behavioral time-courses, showing the performance for each FP duration (25 different durations), separately for each distribution condition. Lines: group mean, shaded area: 95% CI (bootstrapping, 1000 iterations). Both (**c**) accuracy and (**d**) RT show a nuanced relation to FP duration, which is highly dependent on the temporal context.

Notably, the effect of FP duration on performance was strongly modulated by the task temporal structure. To test this statistically, we conducted two one-way ANOVAs (separately for RT and ACC) using the difference in performance between the long and short durations (ΔACC and ΔRT) as our dependent variable, and FP distribution as the between subject factor. Both tests were highly significant (ACC: F(2,139)=14.99, p<10^−6^, η_p_^2^=0.18; RT: F(2,139)=48.27, p<10^−15^, η_p_^2^=0.41). ΔACC was largest in the FLIP-EXP condition, followed by the UNI condition, and the EXP condition (*Fig. 3a*). Tukey-HSD post-hoc tests showed that FLIP-EXP was significantly larger than the other two distributions (both q>3.75, p_Tukey_<0.001), but the difference between UNI and EXP was not significant (q=1.58, p_Tukey_>0.25). The ordering between distribution conditions was similar for RT (*Fig. 3b*), and here post-hoc tests revealed significant differences between all three distributions (all q>3.85, p_Tukey_<0.001). Together, this shows that the temporal structure of the task significantly modulated the effect of time on both perceptual accuracy and reaction times, indicating that temporal anticipation is shaped by the specific expectations formed during the task.

### Uncovering the computation behind temporal anticipation

Next, we turned to investigate the computational mechanisms underlying temporal anticipation, utilizing the full behavioral dynamics, with 25 different FP durations sampled for each distribution (*Fig. 3c-d*). To account for the paucity of data underlying the time-courses of single participants (especially in the low probability durations in the EXP and FLIP-EXP conditions), we focused on prediction of the group averages (see also ref. ^18–20^). Having established in the previous section that the influence of temporal anticipation is deeply tied to the task-level temporal structure, we used the PDF of the FP distribution as the basis of the computation. We constructed multiple model time-courses based on the PDF reflecting distinct hypotheses about the algorithmic and computational steps that this information undergoes, before it is used to modulate perception and action (*Fig. 1*; <u>Methods\Computational Modelling</u>). The following sections are dedicated to examining each of these computational steps in turn.

To assess how well each model captured the behavioral dynamics, we used linear regression (<u>Methods\Model fitting</u>; note this does not indicate that the computation is linear, as the models themselves contained several nonlinear computations). We fit the models separately to the empirical accuracy and to RT time-courses to allow for the possibility of different computations underlying the modulation in each case. The models were constructed to match the dynamics of anticipation, and as higher anticipation is expected to lead to higher accuracy and lower (faster) RTs, we restricted the fits to maintain a positive relation between the model predictions and the accuracy data, and a negative relation between the model predictions and the RT data. For models which did not satisfy this constraint we used a flat line as the model prediction (see <u>Methods\Model fitting</u> for more details). The fitting results for individual distributions were evaluated using adjusted R^2^ (denoted by 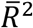) and Akaike Information Criterion (AIC).^26,27^ To evaluate the fits across distributions we used the sum of AIC values from each distribution (see <u>Methods\Model evaluation</u>). Note that models are considered to be a better fit if they have lower AIC. Following the conventions in the literature, models were considered substantially different if they had an AIC difference of more than 4, and were considered slightly different in predicting the data if they had an AIC difference between 2 and 4 (for more information on model comparison see <u>Methods\Model evaluation</u>).^28^

### The HR alone fails to capture the behavioral dynamics of temporal anticipation

As a first step, we examined the two leading candidates for the core computation underlying temporal anticipation: the PDF and the HR. These two temporal functions are tightly related: the PDF is the a-priori probability of S2 appearing at each moment in time, before the trial begins, and the HR is the conditional probability, accounting at each moment for the fact that S2 did not occur yet. The HR can thus be written as PDF/(1-CDF), where CDF is the cumulative distribution function (<u>Methods\Hazard rate</u>), meaning that the PDF serves as the basis for both computations, but the HR adds to it another dynamic transformation (scaling by 1-CDF). As a result, the HR and PDF predict similar anticipation levels for the start of the trial, but as time progresses, they begin to diverge, with the HR predicting growing anticipation relative to the PDF (the HR model may still predict constant or decreasing anticipation over time, if the PDF showed a strong enough decrease, see the EXP condition in *Fig. 4a*).

**Fig. 4:**
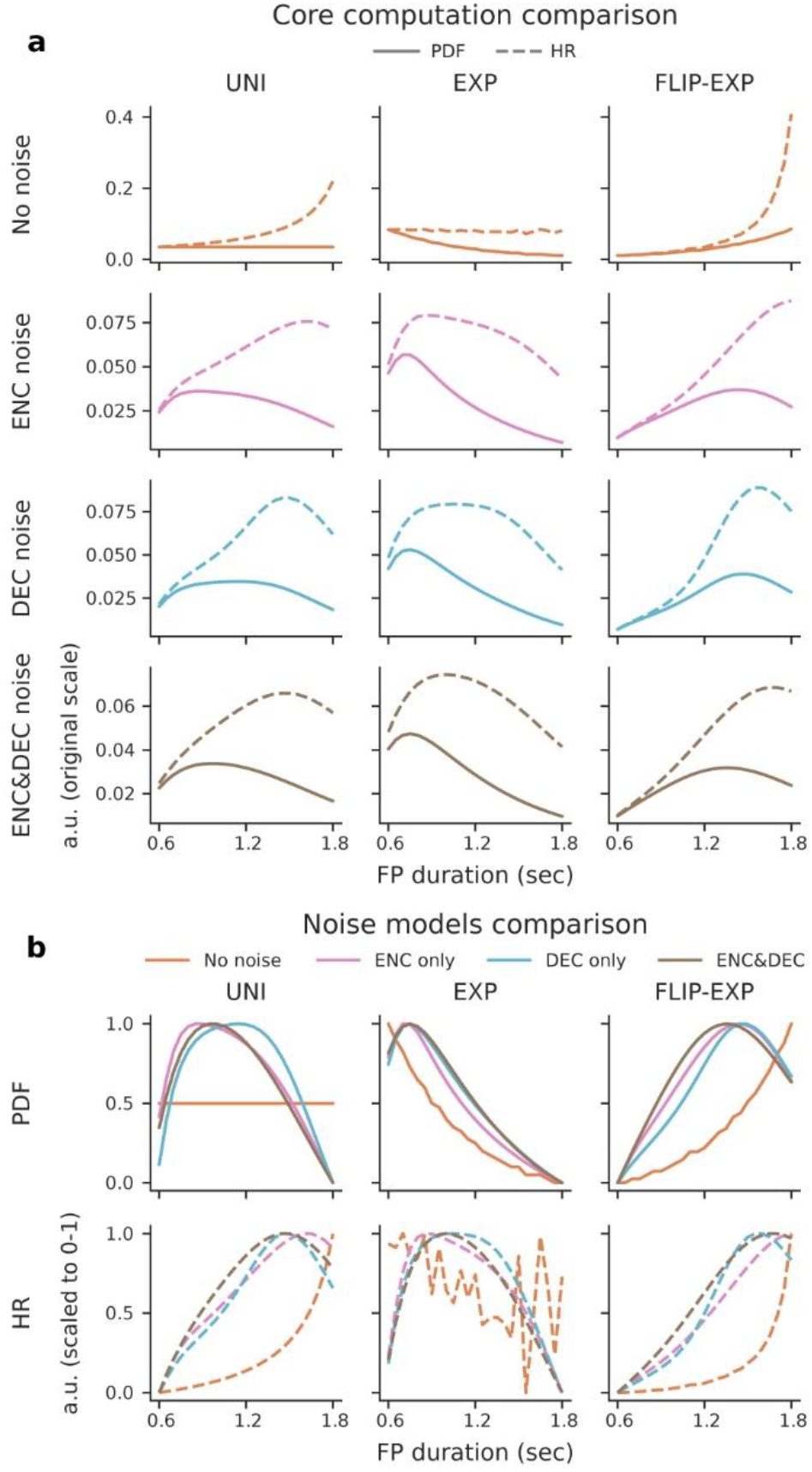
Computational models predicted dynamics. (**a**) Comparison of PDF-based and HR-based models. Solid lines: PDF, dashed lines: HR. Columns: distributions (uniform, exponential, flipped-exponential). Rows and color coding: noise model (noise free, encoding only, decoding only, encoding and decoding; all shown with time-based noise modulation). The PDF and HR models predict similar anticipation levels for short FPs, but they diverge for longer FPs, with the HR predicting higher anticipation relative to the PDF. (**b**) Noise models comparison (time-based noise modulation). Color coding: noise model (as in (a)). Columns: distributions (as in (a)). Rows: core computation (PDF and HR). To ease the comparison between the noise free model and the other models, in this panel all models were scaled to the same range. Compared to their noise free counterparts, models incorporating noise predict low anticipation for the short or long ends of the distribution, especially when the noise free prediction is high. This difference is more pronounced for the long FPs which are noisier under time-based modulation.

To compare between the PDF and HR models, we computed their predictions for each distribution condition (*Fig. 4a* top row) and fit the resulting model dynamics to the empirical ACC and RT time-courses, as explained in the previous section. We focus first on the EXP and FLIP-EXP conditions, which were designed to compare between the PDF and HR models, as PDF_EXP_ and PDF_FLIP-EXP_ are mirror images of each other, but the asymmetric nature of time results in very different trends for HR_EXP_ and HR_FLIP-EXP_. Examining the empirical behavioral dynamics (*Fig. 3c-d*), it is evident that neither the ACC nor the RT dynamics of these conditions are mirror images, suggesting that the PDF model does not capture well the empirical data. However, as we show below, for both ACC and RT the HR model fared even worse than the PDF in predicting the empirical dynamics.

Starting with the FLIP-EXP condition, PDF_FLIP-EXP_ and HR_FLIP-EXP_ both predict higher ACC and faster RTs for longer vs shorter durations, consistent with our previous findings for this condition (*Fig. 3a-b*). However, the full dynamics predicted by both models is markedly different from the dynamics of the empirical data (compare *Fig. 3c-d* to *Fig. 4*). Whereas the empirical ACC and RT time-courses show a decelerating pattern (the changes between time-points become smaller as time progresses), both models predict an accelerating pattern (changes increase with time). Since the acceleration was more prominent in the HR_FLIP-EXP_ model relative to PDF_FLIP-EXP_ model, this results in substantially poorer fits for HR_FLIP-EXP_ relative to the PDF_FLIP-EXP_ for both ACC (AIC_PDF_-AIC_HR_=-16.7; *Fig. 5a*) and RT (AIC_PDF_-AIC_HR_=-12.7; *Fig. 5b*).

**Fig. 5:**
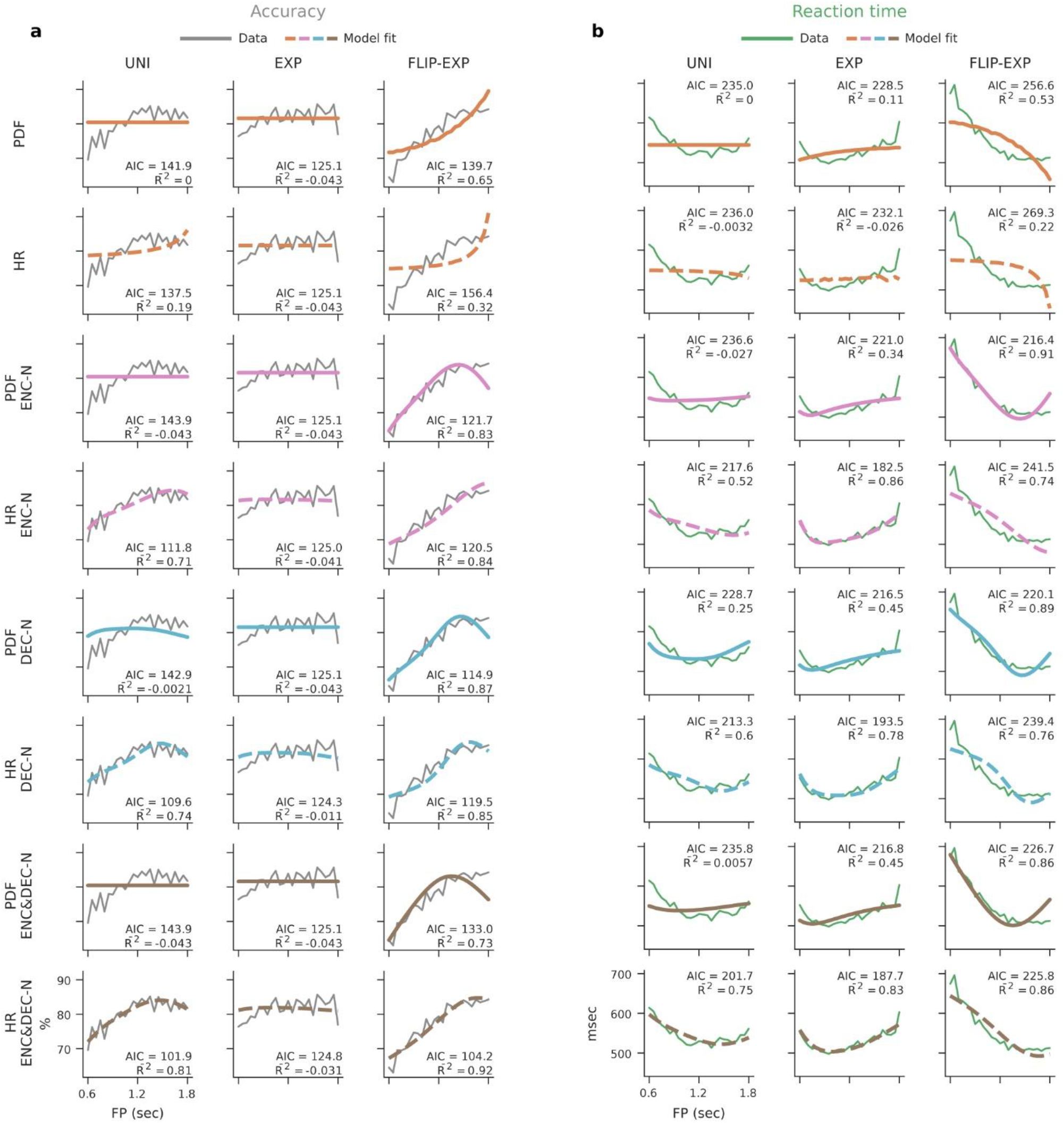
Model fitting results. (**a**) Accuracy and (**b**) RT (correct trials) fitting results. Columns: distributions (uniform, exponential, flipped-exponential), rows: models (solid lines: PDF-based models, dashed lines: HR-based models; colors correspond to the noise model, all using time-based noise modulation, color coding as in *Fig. 4*). Inset text shows the model’s adjusted R^2^ and AIC. Models are considered a better fit if they have higher adjusted R^2^ and lower AIC. Considering only the noise-free models (top 2 rows), the HR model failed to predict the behavioral data. Incorporating temporal estimation noise reversed this pattern and led to better predictions by HR-based models compared to PDF-based models. Overall, the best fit was given by the HR model combining both encoding and decoding noise (bottom row).

Turning to the EXP condition, the empirical ACC time-course was relatively flat, with a small increase after the shortest FPs (*Fig. 3c*). The PDF and HR models both failed to capture this slightly increasing trend, with the PDF_EXP_ model predicting sharp decrease and HR_EXP_ predicting a shallow decrease (see *Fig. 4b* for a scaled version of both models). Therefore, for both models the sign-restricted regression analysis (<u>Methods\Model fitting</u>) led to a flat line fit, and identical AIC values (i.e., AIC_PDF_-AIC_HR_=0; *Fig. 5a*). In contrast to the accuracy data, the empirical RT_EXP_ dynamics are asymmetric U-shaped, with slower RTs for the longest vs shortest FPs (*Fig. 3d*). This slowing at longer FPs is consistent with an overall decrease in anticipation, which is predicted by both PDF_EXP_ and HR_EXP_, therefore both models were able to fit the empirical RT data, with PDF_EXP_ fairing slightly better (AIC_PDF_-AIC_HR_=-3.6; *Fig. 5b*). However, both models did not capture the full dynamics well, in particular, both failed to capture the slower RTs for the short FPs.

Finally, we examine the UNI condition. In this case, PDF_UNI_ predicts a constant level of performance (*Fig. 4*), which does not correspond to our empirical findings (*Fig. 3a-b*). HR_UNI_ in contrast, predicts increasing anticipation with time, i.e., higher ACC and faster RTs. This is indeed what we observed, yet again, the full dynamics are not captured well by the HR model (*Fig. 3c-d*). Nonetheless, compared to the flat prediction of the PDF_UNI_ model, HR_UNI_ fared better in fitting the ACC time-course (AIC_PDF_-AIC_HR_=4.4; *Fig. 5a*), but not the RT time-course, where the AIC values for both models were comparable (AICPDF-AIC_HR_=-1; *Fig. 5b*). Overall, both models did not describe the behavioral dynamics well, however, when considering the results across all distributions, the HR model provided a significantly poorer fit for both ACC (∑(AIC_PDF_-AIC_HR_)=-12.3) and RT (∑(AIC_PDF_-AIC_HR_)=-17.4).

### Computational modelling of temporal-estimation noise

The ‘naïve’ PDF and HR models assume that participants have perfect knowledge of the underlying distribution. This is highly unlikely, as time estimation is inherently noisy.^24,25^ We modelled this by assuming that the subjective FP perceived in each trial is not the objective FP, but a noisy version of it, with additive Gaussian noise (see <u>Methods\Temporal-estimation noise</u>). That is, on trial *i*, with an objective FP of *T*, the subjective perceived duration 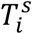 is given by:

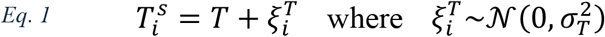

The magnitude of noise in the estimate is determined by *σ*_*T*_, the standard deviation of the noise distribution. We considered two candidates for factors influencing *σ*_*T*_, time-based and probability-based modulation (see <u>Methods\Noise modulation</u>). We focus first on time-based modulation, which assumes estimation noise is proportional to the duration being estimated, in accordance with the ‘scalar property’ of timing (also known as Weber’s fraction in timing).^24,25,29^ Therefore, in this case, we modelled *σ*_*T*_ as a linear function of time, *σ*_*T*_ = *φT* where *φ* is the Weber fraction.

Importantly, we hypothesized that such trial-by-trial temporal-estimation noise can influence both how the representation of the task’s temporal structure is formed and adjusted (how the FP distribution is gradually learned), and the way this information is used on each specific trial to guide behavior. Accordingly, we introduced to our modelling framework two distinct stages (*Fig. 1b*; <u>Methods\Encoding and decoding noise</u>): (A) ‘Encoding’ (forming the temporal representation) – registering the current FP estimate to update the representation of the task’s temporal structure, which forms the basis of the temporal anticipation that modifies responses in future trials, and (B) ‘Decoding’ (reading-out the temporal anticipation) – inferring from the current FP estimate the anticipation in the current trial, which is used to modify the current behavioral response. Both stages may be impacted by noise stemming from the same source. In noisy ‘Encoding’ or ‘Decoding’ the subjective estimated FP is the noisy FP given by Eq. 1, while in noiseless ‘Encoding’ or ‘Decoding’ the subjective estimated FP is identical to the objective FP.

Encoding and decoding as introduced above are both single-trial processes. To enable comparison to the behavioral dynamics (*Fig. 3c-d*), we look at their impact at the task-level, and construct model-predicted dynamics by computing the expected influence (in the probability sense) across all trials. Under this task-level perspective, encoding can be conceived as transforming the external objective PDF to an internal subjective PDF, and decoding as transforming the internal subjective anticipation (which may correspond directly to the subjective PDF or may include additional transformations such as calculation of the HR) to a function representing the anticipation which was effectively used in the task to modulate behavior. Note that the internal subjective anticipation may be represented explicitly in an aggregate form across trials (though this is not essential to our model, see <u>Discussion</u>), yet the task-level decoded anticipation is only a mathematical construct used to compare the model predictions to the behavioral time-courses.

Framed this way, noiseless encoding is equivalent to the identity transformation, mapping the subjective PDF directly onto the objective PDF, and similarly, noiseless decoding is equivalent to an identity transformation between the subjective and objective anticipation. In contrast to this, incorporating the influence of temporal-estimation noise in encoding and decoding results in substantial transformations of the underlying functions, akin to convolving them with a Gaussian kernel. However, the specific input function and the width of this kernel differ between encoding and decoding noise (see <u>Methods\Encoding and decoding noise for the full derivation</u>). Formally:

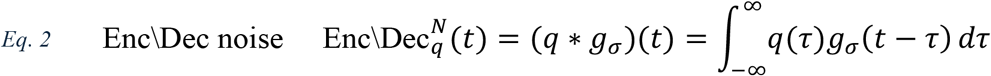

where *q*(*t*) denotes the temporally resolved input function and *g*_*σ*_(*t*) denotes a Gaussian with zero mean and standard deviation of *σ*:

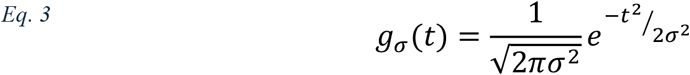

For encoding noise the temporal input function *q* is given by the objective PDF, and the noise level *σ* varies between durations, that is, each objective duration is impacted by its own noise level (*σ*_*enc*_ = *σ*_*τ*_, with *τ* corresponding to the objective PDF *q*(*τ*) in Eq. 3). In contrast to this, for decoding noise *q* is given by the subjective internal anticipation, and the noise level is identical for all durations, determined solely by the decoded duration (*σ*_*dec*_ = *σ*_*t*_, with *t* corresponding to the output 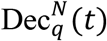 in Eq. 3). See <u>Methods\Encoding and decoding noise</u> for a more detailed explanation.

To gain insight into the impact of encoding noise and decoding noise on the computation, consider the example in *Fig. 1c*, which illustrates their influence on the UNI distribution: First, encoding transforms the objective PDF (panel I) to the subjectively encoded PDF (panel II). In this example encoding is noisy, therefore the perceived FP in every trial may be shorter or longer than the objective FP, resulting in a broader subjective PDF than the objective one, due to ‘sharing’ of the objective probability of each FP duration with the surrounding FPs. When the noise magnitude is higher, this ‘sharing’ is more pronounced. Under time-based noise modulation (where longer FPs are more impacted by noise), this leads to an asymmetric encoded distribution despite the symmetry of the original PDF. Next, we see the influence of internal transformations on the subjective PDF, including the HR computation (panel III) and log-transforming the HR (panel IV), which will be addressed at a later section. Finally, decoding transforms us from the internal anticipation back to the objective FPs, showing the mean anticipation used in responses to each FP duration (panel V). Noisy decoding means that the anticipation used for each trial may correspond to a different duration than the objective FP, and therefore, the decoded anticipation combines the anticipation of the current FP and the surrounding FPs, effectively smoothing the internal anticipation.

### An HR model incorporating encoding and decoding noise best explains behavioral dynamics

As mentioned above, encoding and decoding are part of all models, but not all models incorporate noise in these stages. This results in four models for each of the core computations (PDF and HR) considered previously: (1) Noise free (naïve PDF and HR models) (2) Only Encoding noise (ENC-N) (3) Only Decoding noise (DEC-N) (4) Encoding and decoding noise (ENC&DEC-N). To enable comparison between these models, *Fig. 4b* depicts the predicted dynamics of all four models using the same axes (separately for PDF and HR-based models). This shows that models incorporating noise tend to under-weigh the short or long ends of the distribution compared to the naïve (noise free) PDF and HR models, especially when the naïve model predicts high anticipation. As a result, the PDF models including noise (*Fig. 4b* top) all predict relatively low anticipation for the long FPs, and thus poor performance (low accuracy and slow RT). This trend is (at least partially) mitigated for the HR noise models (*Fig. 4b* bottom), since the HR computation increases anticipation for long FPs.

Behaviorally, we observed improved performance with time in all but one case (*Fig. 3*), which suggests that the HR noise models will outperform the PDF noise models. This is confirmed by the formal model fitting results. Examining first the ACC fits (*Fig. 5a*), the HR noise models substantially outperformed the PDF noise models in the UNI condition (AIC_HR_-AIC_PDF_: −42 to −32.1, comparing each HR model to the PDF counterpart), with the best fit obtained by the model incorporating both encoding and decoding noise (HR-ENC&DEC-N; ΔAIC ≤ −7.7 relative to all other models; *Fig. 5a* bottom row). This same model also substantially outperformed all other models in predicting the FLIP-EXP condition (all ΔAIC ≤ −10.7), while for the EXP condition, where the ACC time-course is nearly flat, no model significantly outperformed a flat line prediction. Combining across conditions (*Fig. 6a*), the best fit for ACC was given by the HR-ENC&DEC-N model (∑ΔAIC ≤ −22.5 relative to all other models), followed by the two other HR noise models (HR-ENC-N and HR-DEC-N), which all substantially outperformed the PDF-based counterparts (*Fig. 6b*; AIC_HR_-AIC_PDF_: −71.1 to −29.5).

**Fig. 6:**
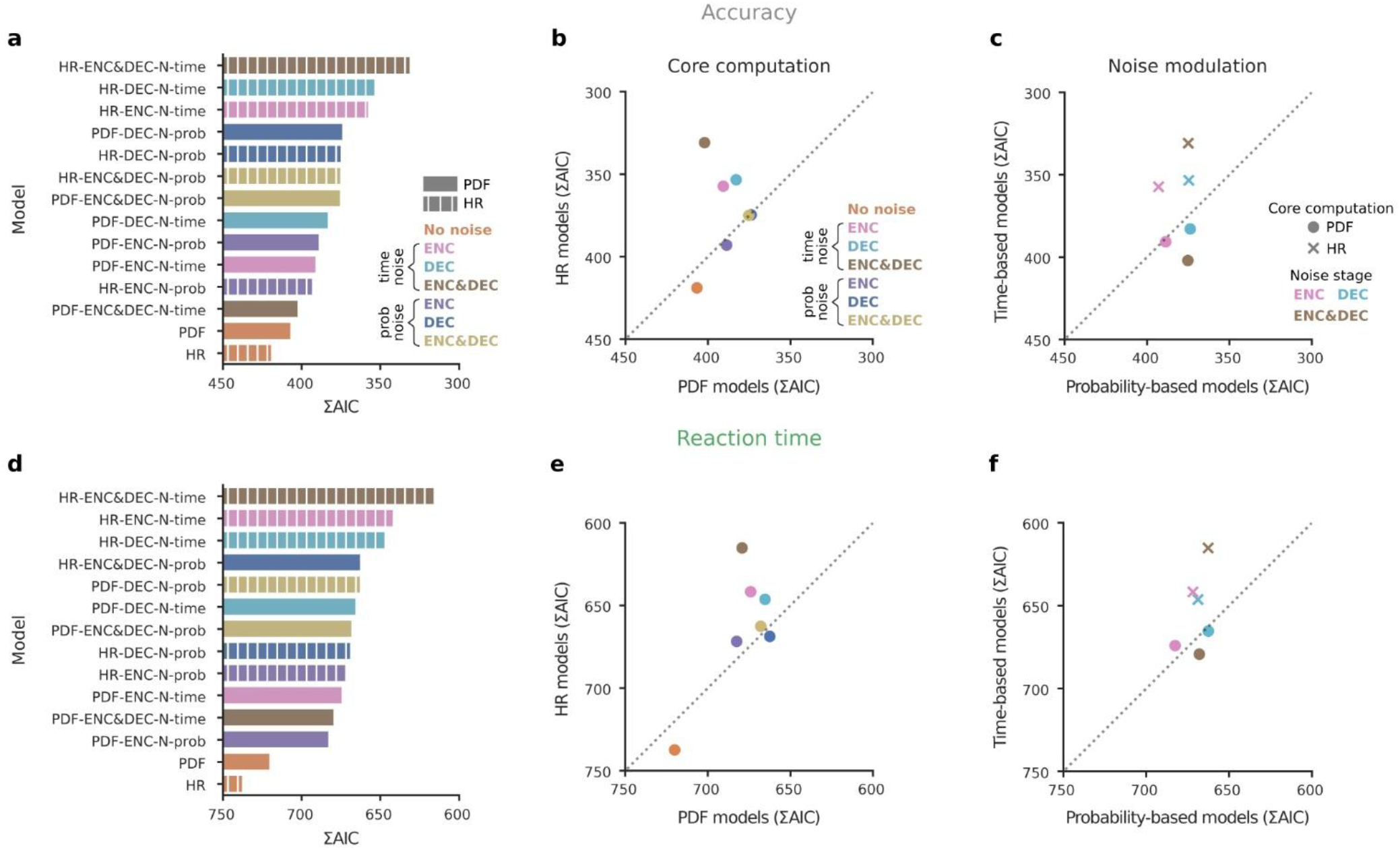
Time-based and probability-based modulation of temporal estimation noise. (**a**-**c**) Accuracy and (**d**-**f**) RT (correct trials) combined fitting results. (**a, d**) Fitting results (∑AIC across distributions) for the noise-free, time-based and probability-based noise models (14 models). Model names indicate the core computation (PDF vs HR), which stage included noise (encoding only (‘ENC-N’), decoding only (‘DEC-N’), or both (‘ENC&DEC-N’) and whether the noise time-based (‘time’) or probability-based (‘prob’). X-axis is flipped so that lower AIC (corresponding to improved fits) is marked by longer bars. For both accuracy and RT, the best fit was given by the HR model with encoding and decoding time-based noise modulation. (**b, e**) Comparison of PDF and HR models. Each dot corresponds to one noise model (color coded). x-value: ∑AIC of the PDF model, y-value: ∑AIC of the HR model (all other modelling steps are identical). Dashed line: identity. Since lower AIC indicates a better fit to the data, axes were flipped, so that dots above the identity line correspond to cases where the HR-based model provided a better fit to the data, and dots below the line correspond to cases where the PDF-based model provided a better fit. For both accuracy and RT, HR-based models outperform PDF-models for time-based modulation, which showed the best fit overall. (**c, f**) Comparison of time-based and probability-based noise modulation (excluding the two noise-free models). Each dot corresponds to one choice of modelling steps (marker color: ENC-N, DEC-N, or ENC&DEC-N; marker shape: PDF or HR). X-value: ∑AIC of the probability-based noise model, y-value: ∑AIC of the time-based noise model. Dashed line: identity. Axes were flipped as in (b, e). For both accuracy and RT, time-based noise modulation outperformed probability-based noise modulation in HR models, which showed the best fits overall.

The RT model fits showed a similar direction (*Fig. 5b*). For the UNI condition the HR noise models substantially outperformed the PDF models (ΔAIC: −34.1 to −15.4), and the best model was again HR-ENC&DEC-N (ΔAIC ≤ −11.6 relative to all other models). For the EXP condition HR models also outperformed the PDF models (ΔAIC: −38.5 to −23), though here the best model was HR-ENC-N (AIC_HR-ENC-NOISE_-AIC_HR-ENC&DEC-NOISE_=-5.1). Finally, for the FLIP-EXP condition neither the PDF nor the HR models fully capture the flattening of the empirical RT curve in the longer FP durations, but the PDF models provided better fits overall. Across conditions, the RT results (*Fig. 6d*) are highly consistent with the ACC results, with the best fit given by the HR-ENC&DEC-N model (∑ΔAIC ≤ −26.5 relative to all other models), followed by again by the other two HR noise models, all substantially outperforming their PDF counterparts (*Fig. 6e*; AIC_HR_-AIC_PDF_: −64 to −19.1).

Together, these results show that considering temporal-estimation noise is crucial to understand the computation underlying temporal anticipation. Although the naïve HR model did not predict the data well, incorporating time-based modulated temporal-estimation noise in the computation substantially improved the fit, and led the HR-based models to substantially outperform the PDF-based models (*Fig. 6b,e*). Moreover, we found that the best prediction was given by the HR-based model incorporating both encoding and decoding noise, suggesting that estimation noise influences both the formation of the temporal representation and how this representation is used to guide behavior.

### Time-based modulation of estimation noise provides a better fit to behavior compared to probability-based modulation

As an alternative to time-based modulation of estimation noise we considered probability-based modulation (*Fig. 1a*), promoted recently by Grabenhorst and colleagues.^18–20^ Under this hypothesis, estimation noise is not modulated by the FP duration itself, as is assumed by time-based modulation, but by the probability of encountering each FP duration. Thus, in this case we model estimation noise is a linear function of probability, i.e., *σ*_*T*_ ∝ *f*(*T*), with higher probability leading to *lower* noise levels (<u>Methods\Noise modulation</u>). Such a reduction can stem, for example, from learning effects following the increased exposure to high probability FPs.^30^

We computed the model predictions for each probability-based noise model, fit them to the empirical accuracy and RT dynamics, and compared the fits to those obtained by models incorporating time-based noise modulation. HR models incorporating probability-based noise modulation were substantially inferior then the time-based noise models in accounting for the trends of the behavioral data for both ACC (all AIC_HR-PROB_-AIC_HR-TIME_ ≥ 21.3, *Fig. 6c*) and RT (all AIC_HR_-PROB-AIC_HR_-TIME ≥ 22.4, *Fig. 6f*). This was not the case for PDF noise models, which showed a more nuanced picture, and even some improvement under probability-based modulation. Yet, since HR models generally outperformed the PDF based models (*Fig. 6b,e*), when we consider all models together the best fit for both ACC and RT was obtained by a time-based model (HR-ENC&DEC-N), which outperformed all probability-based models by a substantial margin (ACC: ∑ΔAIC ≤ −42.9, *Fig. 6a*; RT: ∑ΔAIC ≤ −47.3, *Fig. 6d*), followed by the other two time-based HR noise models which also significantly outperformed all probability-based models (ACC: ∑ΔAIC ≤ −16.5, RT: ∑ΔAIC ≤ −16.2). To conclude, we found that for both ACC and RT probability-based noise models were less accurate in predicting the behavioral dynamics compared to time-based noise models.

### Logarithmic transformation of the HR best captures the computation underlying temporal anticipation

As a final step, we considered the possibility that the cognitive representation of temporal anticipation involves additional nonlinear transformations beyond the HR. Specifically, we considered three such transformations: (1) Logarithmic (2) Reciprocal (1/*p*) and (3) Exponential (*Fig. 1a*; <u>Methods\Nonlinear transformations</u>). Logarithmic transformation was considered because it was shown to underly neural coding schemes across domains^31,32^ and it features in core explanations of psychophysical phenomena.^33–35^ Moreover, the logarithmic transformation has theoretical significance in the context of Shannon’s information theory, where the information content (or ‘surprisal’) of an event with probability *p* is defined as −*log*(*p*).^36^ Thus, incorporating a logarithm in the computation implies responses are not influenced by the raw probability of an event (be it the a-priori probability, given by the PDF, or the conditional probability, given by the HR), but by how surprising it is. In effect, this accentuates the impact of low-probability events, predicting even larger behavioral differences between short and long FPs compared to models using the raw PDF or HR. Reciprocal transformation was included because it was supported by recent work on temporal anticipation.^18–20^ Conceptually, this transformation implies that responses are not related to the probability of each FP, but to the number of trials one needs to wait for in order to observe that duration. This results in an even larger accentuation of low probability events (those requiring a longer wait) than that predicted by the logarithmic model. Finally, exponential transformation was considered for theoretical completeness, because it is mathematically complementary to the logarithm, and therefore has the opposite effect on predictions.

We incorporated the transformations into our modelling framework by applying them to the internal representation prior to decoding (*Fig. 1b*). Each transformation was evaluated in combination with both the PDF and HR computations, and all noise models (including the noise-free model and both time-based and probability-based noise models). Overall, this resulted in 14 models per transformation and 56 models in total (including 14 models without a nonlinear transformation), which were all fit to the accuracy and RT time-courses of each distribution condition.

Comparing the influence of nonlinearities across all models, we found that in most cases incorporating the logarithmic and reciprocal transformations significantly improved the model fits compared to the model without a nonlinear transformation (except for the models including only decoding noise; *Fig. 7a-b*). In contrast, the exponential transformation rarely improved model fits relative to the model without a nonlinearity. Comparison between the logarithmic and reciprocal transformations was more nuanced. Although there was a tendency for the reciprocal transformation to provide better fits than the logarithmic transformation, this was driven by models with overall lower rather than higher correspondence to the behavioral data (poorer fits). That is, reciprocal transformation improved the performance of otherwise weaker models, but the best fitting model for both behavioral measures was based on a logarithmic transformation. Specifically, for both ACC and RT, the best fit to the behavioral time-courses was provided by the model based on the HR followed by a logarithmic transformation, and incorporating time-based encoding and decoding noise (HR-log-ENC&DEC-N-time; ∑ΔAIC relative to all other models, ACC: ∑ΔAIC ≤ −7.1, *Fig. 7c*; RT: ∑ΔAIC ≤ −12.1, *Fig. 7d*).

**Fig. 7:**
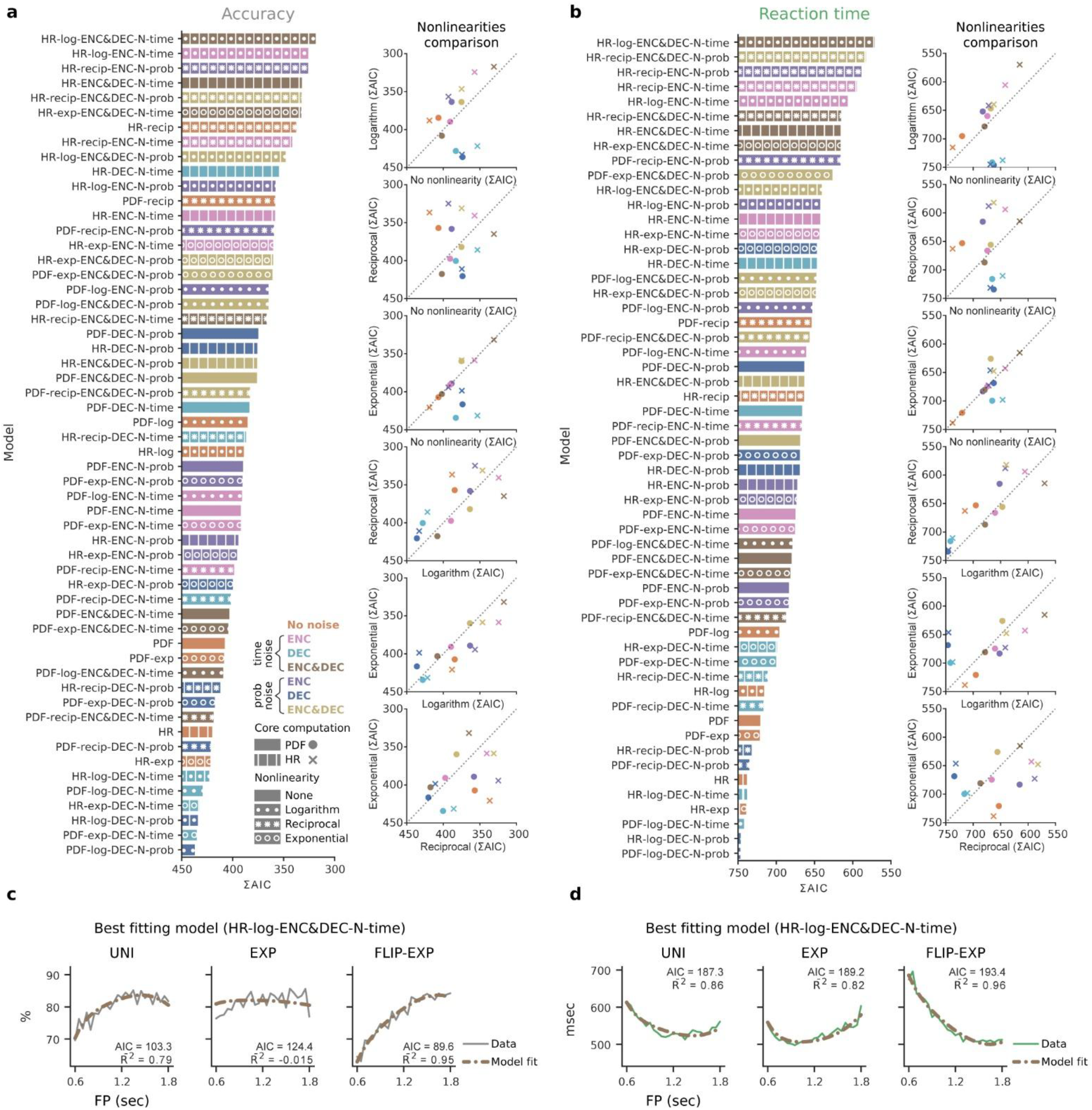
Effect of nonlinear transformations. (**a**) Accuracy and (**b**) RT (correct trials) combined fitting results (∑AIC across distributions) for all 52 models (14 without any nonlinearity (*Fig. 6*) and 14 for each of the three nonlinearities: logarithm, reciprocal and exponential). X-axis is flipped so that lower AIC (corresponding to better fits) are marked by longer bars. Scatter plots: comparisons between nonlinearity models, and to absent nonlinearity (each panel compares two cases). Single dots: one base model (marker shape: PDF or HR, marker color: noise model). Dashed line: identity. Here too, axes were flipped, so that dots above the identity line indicate that the y-axis nonlinearity provided a better fit to the data, and dots below the line indicate that the x-axis nonlinearity provided a better fit. (**c**) Accuracy and (**d**) RT (correct trials) best model fitting results. Columns: distributions (uniform, exponential, flipped-exponential). In both cases, the best fit was given by the HR-based model with logarithmic nonlinearity, incorporating time-based noise modulation in both the encoding and decoding stages.

Finally, incorporating nonlinear transformations into the computation did not alter our previous conclusions (*Fig. 7a-b*). In particular, HR-based models still substantially outperformed PDF-based models both when considering all models together (ACC: 10 of the 10 best fitting models were HR-based, RT: 8 of the 10 best fitting models were HR-based), and for each nonlinear transformation separately. Additionally, for HR models time-based noise modulation continued to outperform probability-based modulation. These results support the robustness of our core findings and highlight the importance of nonlinear scaling for capturing of the internal computation underlying temporal anticipation.

## Discussion

In a world with constant changes, being able to anticipate the time of events offers substantial evolutionary advantages, as it allows organisms to prepare responses in advance and to allocate cognitive and energetic resources effectively at the time when changes are expected to happen.^2,3^ Environmental temporal structures provide a critical source of information which organisms can draw upon to form anticipatory predictions. In this study, we examined the influence temporal anticipation has on behavior, how anticipation is modulated by temporal structures, and the computational processes behind this modulation. First, we show that temporal anticipation not only facilitates motor response but also perceptual processing, and that both effects are modulated by the temporal structure. Second, using computational modelling we show that the internal computation behind the modulation is best approximated by the logarithm of the event hazard rate, while also revealing a key role for temporal-estimation noise both at the encoding stage, when the temporal representation is formed, and at the decoding stage, when it is used to guide behavior.

To uncover these findings, we employed a paradigm combining change discrimination with variable FP. The task was maintained at a perceptually challenging threshold using an ongoing staircase procedure. The merit of this design lies in the fact that it allowed us to simultaneously measure the influence of temporal structure on perceptual accuracy and reaction times, providing a comprehensive view of how temporal anticipation affects behavior. Importantly, we manipulated the FP distributions across subjects rather than within subjects across different blocks, as is often done in similar studies (e.g., ref. ^17–20^). This design choice minimized carryover effects between distributions, ensuring that participants’ behavior reflected adaptation to a single temporal structure, rather than contamination from prior learning. Moreover, we used a dense temporal sampling, with 25 different FP durations for each distribution condition, which enabled us to dissect the underlying computation in greater detail than previous studies, many of which employed only a handful of FP durations (e.g., ref. ^10,14,16,37,38^).

### Temporal structure modulates perception and action

Prior studies examining distribution-driven temporal anticipation have predominantly focused on how it influences reaction times.^7,8,10,11,13,15–20^ Effects on response speed that were identified by these studies were traditionally attributed to non-specific motor preparation, as the preparatory signal marking the start of the anticipatory period does not contain information about the target.^7,15,16^ This interpretation was further supported by studies showing that FP manipulations influence other variables that are closely tied to the motor system such as the force of the response and evoked motor potentials (for an overview of these findings, see ref. ^39^). Our finding of strong modulation of perceptual acuity by distributional temporal structure challenges this motor-centric view and suggests broader role of distribution-driven temporal anticipation in shaping cognition.

This conclusion is also in agreement with studies which manipulated temporal anticipation through other means, such as cued associations (when a cue indicates the duration of the FP) or rhythmic patterns, and have reported influences of temporal anticipation on perceptual abilities (for example ref. ^40–42^, though see ref. ^43^). Generalizing a-priori from these forms of anticipation to distributional temporal information is unwarranted, as different forms of temporal information may exert their influence through distinct cognitive and neural mechanisms (e.g. top-down modulation vs more local mechanisms such as neural entrainment).^3,44^ Nonetheless, our results suggest a deeper connection between different forms of temporal anticipation, and support the view that temporal predictions exert diverse influences across perceptual, cognitive, and motor domains.^2,45^

Additionally, showing that perceptual acuity is modulated by temporal information has implications for the neural correlates of temporal anticipation. Prior work has shown that anticipatory signals are present in many cortical areas, including both control regions, such as LIP and premotor regions, as well as sensory regions.^8,12,13,20,38,46,47^ Whereas most of these studies examined temporal expectations in isolation, there is also some evidence that temporal information can interact with coding of other types of information. For example, it was shown that in visual cortex, temporal anticipation modulated neural selectivity for spatial location.^22,48^ Based on our findings, we predict that temporal anticipation will also interact with the coding of specific visual features, for example by improving orientation tuning at anticipated moments of change, yet this remains to be shown.

### The computations underlying temporal anticipation

To uncover the computational and cognitive processes underlying the influence of environmental temporal structure on behavior, we constructed multiple computational models, reflecting a broad range of hypotheses on how temporal predictions are processed. Each model was evaluated for its ability to predict the observed behavioral dynamics for three FP distributions and was contrasted with other models both quantitatively and qualitatively. Using this approach, we revealed that the internal computation is best captured by the logarithm of the event HR, and that this computation is further shaped by temporal-estimation noise at both the encoding and decoding stages. Thus, our results show that temporal anticipation arises from an intricate interplay between external environmental structure and internal processing limitations.

### The hazard rate and the Bayesian perspective

The HR represents the conditional probability of an event occurring at each moment, given that it has not occurred yet. Returning to the bus example introduced earlier, the HR embodies the rationale that as time progresses and the bus has yet to arrive, our expectation for its arrival should intensify relative to the a-priori expectation (provided that the overall likelihood of arrival was high to begin with, i.e., a low or moderate ‘catch rate’). The HR computation is not only intuitive, but also optimal computationally, exemplifying the Bayesian approach to updating prior expectations in light of evolving evidence (which is, in this case, the ongoing absence of the anticipated event).

Despite its theoretical appeal, the HR is computationally demanding and potentially unstable, as it requires dividing the PDF by the complement of the cumulative distribution function (1-CDF), a value that often approaches zero. Thus, while many studies have suggested that the HR underlies temporal predictions,^2,3,8,13,14,21,22,49^ these conclusions were often drawn without explicitly comparing alternative models or relied on data limited to a few foreperiod durations, leaving the robustness of the HR account uncertain. Moreover, some recent studies failed to find evidence for the HR in temporal anticipation (e.g., ^47^). In particular, recent work by Grabenhorst and colleagues^18–20^ suggested that temporal implementation may instead be based on a simpler computation, the reciprocal of the event PDF. Our findings do not support their conclusion. Instead, we show the HR is crucial for capturing the dynamics of behavioral influences of temporal anticipation. Thus, our findings imply that temporal anticipation is, in a sense, Bayes optimal, situating time perception within the broader framework of the “Bayesian brain” hypothesis, which has been highly successful in explaining various aspects of neuroscience and psychology.^50,51^ However, our account also deviates from strict optimality in two important ways, temporal-estimation noise and logarithmic nonlinearity, which will be discussed below (see ref. ^52^ for other cases of suboptimality in decision making).

### The role of temporal-estimation noise

We developed a computational framework to understand the influence of temporal-estimation noise on performance, examining separately the influence on encoding of temporal structures and on decoding this information on a trial-by-trial basis. Mathematically, this manifests as two related but distinct algorithmic steps: encoding noise influences the subjective representation of the PDF while decoding noise influences the way temporal information is used to guide behavior. Related mathematical operations have been applied in prior literature (e.g., ^8,48^), with some studies smoothing the objective PDF prior to computing the HR (mimicking the influence of encoding noise, albeit imprecisely), and others smoothing the HR itself, after it was computed with the objective PDF (mimicking the influence of decoding noise). However, to the best of our knowledge, no studies have accounted for both noise manifestations, which we show is needed to capture accurately the behavioral dynamics, as the model incorporating both noise processes outperformed models that considered either one in isolation. Moreover, our work situates both operations within a single unified framework, revealing that both derive naturally from a single uncontroversial assumption: that subjective temporal estimates deviate from objective durations due to inherent variability in time estimation. Thus, our unified perspective not only improves the accuracy of behavioral predictions but also provides a more comprehensive understanding of the interplay between processing limitations and internal inference in orchestrating adaptive behavior.

We considered two primary hypotheses regarding the underlying source of estimation noise: time-based modulation and probability-based modulation. Time-based modulation is based on the scalar property of time, which posits that variability in time estimation increases proportionally with the interval being estimated. This principle has been widely supported in the timing literature,^24,25,29^ but has also been shown to vary between different time scales and contexts.^53,54^ By contrast, probability-based modulation, as proposed by Grabenhorst and colleagues,^18–20^ suggests that noise levels depend on the probability of encountering specific durations, and specifically, that more probable durations are associated with reduced noise, potentially due to learning effects following greater exposure.^30^ In our hands, time-based noise modulation provided a substantially better fit to the data compared to probability-based modulation. This supports the relevance of the scalar-property for understanding temporal anticipation. However, further work is needed to determine whether it also contributes to temporal anticipation across broader FP ranges and other contexts.

### Logarithmic scaling and efficient information coding

Beyond the HR, we show that incorporating into the computation a nonlinear transformation which emphasizes lower probabilities substantially improves the model fits to both accuracy and RTs. For this role we considered the logarithmic and the reciprocal transformations. Both fared well, with the best fit being obtained by the logarithmic transformation. This finding aligns well with Shannon information theory, where the logarithm of probability (known as Shannon surprisal or Shannon information) quantifies the ‘unexpectedness’ of an event.^36^ Thus, the log-transformed HR can be interpreted as encoding the informational weight of an event, making it a natural candidate for modulating behavioral responses to anticipated events. Moreover, logarithmic transformations feature broadly in psychophysical laws and in neural coding schemes. For example, Fechner’s law describes how perceptions of stimulus intensity (e.g., brightness or loudness) scale logarithmically with physical intensity,^33^ and in the context of neural representations, logarithmic relationships have been proposed to explain sensory coding of various quantities (e.g., ref. ^31,32,35^). Our findings extend these principles to temporal anticipation, implying that the brain uses logarithmic transformations, or a similar concave scaling operation, as a general mechanism across domains to compress large numerical ranges and optimize coding efficiency.

### Constraints on mechanistic modelling

While our computational account provides important constraints on the underlying neural mechanisms, it does not fully determine them. Specifically, our finding of a shared computational explanation for the influence of temporal anticipation on perception and action should not be taken to imply that both rely on a single centralized mechanism. The observed effects may be the result of a single top-down modulatory source but can also emerge from more localized processes executing similar computations in different brain regions. Additionally, although the noisy HR-logarithm model captures well the behavioral dynamics, this does not indicate that these exact transformations are explicitly represented within the brain; for example, that the brain first computes the HR and then applies a logarithmic transformation. Instead, the brain may approximate the result of both computations together. Moreover, this approximation may arise indirectly from other mechanisms or processes. For example, the multiple trace theory of temporal preparation (MTP) suggests that the behavioral manifestations of temporal anticipation arise from memory traces formed by exposure to previous trials,^17,23^ which can take a form that is similar to our proposed computations, without carrying them out sequentially.

Finally, we do not mean to imply that the anticipation for all FP durations is necessarily represented simultaneously in the brain, nor that the brain ‘reads out’ an explicit level of anticipation at any given moment. While this could be the case, the implementation could also rely on a sequential process, such as a neural population traversing through a series of states where each state of the neural trajectory corresponds to a specific FP duration (see ref. ^55^ for the importance of such coding mechanisms in interval timing, and ref. ^47^ for a specific example pertaining to temporal anticipation). Under this interpretation, encoding could be reflected in the overall structure of the trajectory,^56,57^ while decoding may depend on the specific state of the neural population at the moment the event occurs.^58^ Thus, encoding noise may manifest as deviations in the structure of the trajectory due to inaccuracies in the formation, and decoding noise could emerge from variability in the speed at which the neural state progresses through its trajectory, leading to discrepancies between the anticipated and actual timing of events.

We do not resolve between these options. Instead, our findings provide constraints on mechanistic modelling, which must be capable of carrying out the computations specified by our model. Future studies integrating modelling with neuroimaging or electrophysiological data will be essential to elucidate the mechanisms by which these computations are executed and to bridge the gap between algorithmic and neural descriptions.

### Limitations and future directions

First, while our task design offered several advantages, in particular, the ability to measure the influence of anticipation on both perceptual accuracy and reaction times, it inherently combines elements of temporal anticipation with temporal attention, which some authors have suggested are distinct processes with potentially different influences on behavior.^59,60^ Future work is needed to dissociate these factors, for instance, by systematically varying the attentional demands of the task while holding the temporal structure constant. Moreover, the influence of task-specific factors, such as reward contingencies or motivation levels, on how temporal anticipation exerts its effects, remains to be tested.

Second, while our computational model provided a good fit to the aggregate behavioral data, it does not capture trial-by-trial variation. Such work can examine how the probability distribution of different intervals is learned over the course of the experiment, and through this also inform our understanding of encoding noise. Modelling trial-by-trial variation can also lead to insights regarding sequential effects in temporal anticipation, such as the influence of the FP of the previous trial on the current trial performance.^10,16,61^ Additionally, future work can extend our account of noise in the computation, for example, by understanding the role of noise in estimating the probability of no event occurring (the ‘no-change’ rate). Finally, as mentioned above, our model does not address the mechanistic implementation of the computations, which will require combining computational modelling with neural recordings.

## Conclusions

This study addressed two key questions: first, whether the influence of temporal structure extends beyond motor responses to shape perceptual acuity, and second, what computational mechanisms underlie this process. To answer the first question, we used a novel paradigm integrating a challenging change discrimination task with variable foreperiods and show that temporal anticipation facilitates both perceptual and motor processing. Next, we used computational modelling to shed light on the cognitive operations behind these influences, revealing that they are best captured by a logarithmic transformation of the hazard rate, and that temporal-estimation noise plays a pivotal role in shaping the computation. These findings contribute to a growing understanding of how the brain extracts and leverages temporal information to guide adaptive behavior, providing insights into the interplay between environmental regularities and internal processing constraints in this process. Together, this work provides a comprehensive framework for understanding the cognitive computations underlying temporal anticipation and their role in shaping perception and action.

## Methods

### Participants

185 participants completed the task. Participants were of both sexes (95 female, 90 male), between 18 and 45 years old (mean ± SEM: 27.6±0.5 years). Recruitment was done through Prolific (https://www.prolific.com/). Participants received 9$ for their participation (∼12$ per hour). Participant exclusion criteria were: 1. Not having normal or corrected to normal vision, 2. Currently taking any medication to treat symptoms of depression, anxiety or low-mood, 3. Being diagnosed with mild cognitive impairment or dementia, 4. Considering themselves to be neurodivergent and 5. Considering themselves to have attention deficit disorder (ADD) or attention deficit hyperactivity disorder (ADHD). These criteria were assessed by self-report questions prior to participation. Participants were told that the study investigates our ability to identify and discriminate visual changes but were naïve regarding the timing aspect of the study. Ethical approval was provided by the institutional ethics committee at the Hebrew University of Jerusalem and all participants have given their informed consent to participate.

### Assignment to conditions

Participants were assigned an FP distribution condition based on the order of entering the experiment website. Participants were first directed to Morys-Carter’s website^62^ which assigns participants consecutive numbers based on the order of their entry, then, based on this number they were divided into conditions in alternation. Since some participants aborted the task after being assigned a number this resulted in uneven distribution of participants to the different conditions (N_UNI_=58, N_EXP_=59, N_FLIP-EXP_=68).

### Participant exclusion

#### Setup

We excluded 16 participants whose frame rates were identified by the experiment code to be below 45 Hz, suggesting inconsistent frame rates or unusual setup. Additionally, participants were instructed to perform the experiment on a laptop or desktop computer, however, one participant indicated using a 3.3-inch device (diagonal, 1.6 by 2.9 inch) for performing the task and was therefore excluded (all other participants indicated screen sizes were above 10-inch, in agreement with the task requirements instructions; see <u>Setup calibration</u> regarding measurement of screen sizes).

#### Performance

The task included a staircase procedure designed to ensure ∼79.5% accuracy across the experiment (more details under <u>Task</u>). The vast majority of participants reached accuracies very close to this desired value (170/185 of participants were between 78-81%). We excluded participants with accuracy that was more than 5% of this value, since it indicated the task was not performed properly in these cases (this resulted in 9 additional excluded participants: 8 performed poorly below 74.5%, and one above 84.5%). Note that all of the excluded participants reached either the minimal or maximal values of possible change magnitude sizes (which were very wide, see <u>Staircase procedure</u>), which serves as additional confirmation that they were either not performing the task, or the task was not well adapted to their setup or their perceptual limits.

### Attention checks

To ensure that we were only considering participants who were attentive throughout the task, we excluded all participants that reached a change magnitude larger than 14° during the task (5 steps above the starting value in our logarithmic scale staircase procedure, see <u>Staircase procedure</u>) or had 12 or more trials with out-of-time responses throughout the task (resulting in another break in the task; see <u>Repeated trials</u>). This led us to exclude 17 additional participants (7 participants who reached the maximal change magnitude of 39.1°, 7 who reached change magnitudes of 14.4-39°, and 3 more participants who had a large number of out-of-time trials).

In total, this left N_TOTAL_=142 subjects (67 female, ages (mean ± SEM): 27.4±0.6 years), across the three conditions: N_UNI_=47 (22 female, ages 28.1±1 years), N_EXP_=41 (23 female, ages: 28.4±1.1 years), N_FLIP-EXP_=54 (22 female, ages 26.1±0.8 years).

## Task

### Trial procedure

Participants performed a novel task, combining visual change discrimination with temporal anticipation, analogously to traditional variable foreperiod (FP) designs.^7^ The task included two types of trials: 1. Standard (change) trials (∼87.5%): participants were presented with two consecutive oriented gratings and required to judge the direction of orientation change between them (clockwise or counterclockwise). 2. No-change trials (∼12.5%): only one grating was presented, without any change, and no response was required. All trials began with the presentation of a central fixation cross for 0.5-1 sec (discretized at 0.05 sec steps, uniform distribution across trials). These were followed by trial-type specific routines:

#### Standard (change) trials (*Fig. 2a*)

Two tilted gratings were presented sequentially (with a central aperture displaying the fixation cross over the gray background; see <u>Stimuli</u>): S1 was presented for 0.6-1.8 sec, drawing from one of three distributions (between participants manipulation; <u>Foreperiod distributions</u>; *Fig. 2b*), followed by S2 presented briefly for 0.05 sec.

In this sequential design the onset of S1 serves as a preparatory cue, providing participants with probabilistic knowledge about the timing of S2, when they can perform the task. Therefore, the duration of S1 serves as the FP duration and the distribution of S1 durations as the FP distribution. S2 was followed by 0.05 sec of only fixation. Then, a response screen appeared displaying the question (‘What was the change?’) and the key-response mappings (‘X’ for counterclockwise and ‘N’ for clockwise rotations; these keys were selected because their location is consistent across the most commonly used Latin-script keyboard layouts worldwide). Responses were only permitted from the onset of this screen. Following the participant’s response feedback was presented for 0.7 sec, which included changing the color of the fixation cross (green\red for correct\incorrect trials respectively), and verbal feedback (‘Correct!’\’Incorrect!’) above the fixation, in matching color.

#### No-change trials

S1 was presented for 1.85 sec (corresponding to the longest S1 duration + S2 duration), without any additional grating. This was followed immediately by a 0.7 sec screen with the words ‘No change’ in blue and a corresponding fixation color change. Participants were instructed to avoid pressing any button during these trials. No-change trials were used as a type of ‘catch trials’, to ensure that participants were waiting for a change to appear before responding, and to guarantee that the hazard rate did not rise to infinity at the end of the range of stimuli durations (see <u>Hazard rate</u>).

### Foreperiod distributions

The duration of S1 was varied on a trial-by-trial basis, drawing from a predefined FP distribution (per participant). We used three distributions: uniform (UNI), exponential (EXP) and flipped-exponential (FLIP-EXP). The EXP and FLIP-EXP distributions were chosen since the distributions themselves are mirror images of each other, but their hazard rates progress in opposite directions (*Fig. 4b*, <u>Hazard rate</u>), enabling us to dissociate between the two central predictions regarding the core computation underlying temporal anticipation. The UNI distribution was chosen to enable easy comparison with the literature (it is used most frequently in many tasks), and since it offers an intermediate step between the EXP and FLIP-EXP distributions with regard to many model predictions (*Fig. 4*).

For all distributions, we used the interval between 0.6 to 1.8 sec, discretized at 0.05 sec steps, resulting in 25 different durations. We aimed to reach a no-change (catch) rate of approximately 12.5%, and therefore the sum of the probability density function (PDF) was adjusted to reach ∼0.875 in all cases.

For the uniform distribution we used the following PDF:

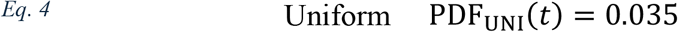

We set the number of trials in each duration to 18 (3 for each S1 orientation, see <u>Stimuli</u>). This resulted in 450 change trials all together, to which we added 64 no-change trials, totaling to 514 trials with a no-change rate of 12.45%.

For the exponential and flipped-exponential distributions we used the following PDFs, which are identical when mirrored around 1.2 sec:

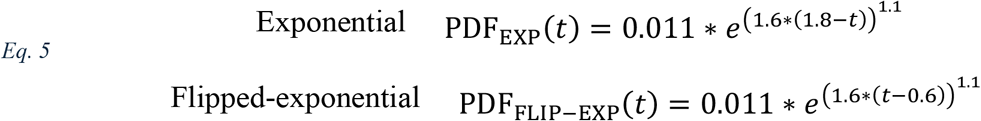

To arrive at the number of trials for each duration we first scaled these PDFs to ensure that all FPs will have a minimum trial count of six. We then performed a modified rounding procedure to arrive at the final number of trials for each duration (the result was similar to standard rounding but resulted in the two distributions deviating slightly from perfect mirror images). This resulted in 482 change trials for each of the two distributions, to which we added 68 no-change trials, reaching a total of 550 trials with a no-change rate of 12.36%.

### Stimuli

All stimuli were presented centrally on a mid-grey background (RGB: 128, 128, 128). Prior to performing the task participants underwent a procedure to measure their screen size and were asked to sit 60 cm away from the screen (see <u>Setup calibration</u>). Stimuli sizes and spatial frequencies reported below in visual angles assume both steps were performed correctly. The fixation cross size was 0.68° x 0.68° visual angles (line width 0.08°). Fixation color was black throughout the task, except for the feedback screen where it was green (correct trials), red (incorrect trials) or blue (no-change trials). The main stimuli (S1 and S2; *Fig. 2a*) were tilted grayscale gratings, displayed at 0.8 contrast, overlayed with an aperture for displaying the fixation cross, which was implemented by applying a custom mask to the gratings. The mask was fully transparent between two concentric circles with diameters of approximately 1.9° visual angles (inner circle) and 4.1° visual angles (outer circle), and then tapered visibility off gradually towards the inside of the circle and to the outside, corresponding to Gaussian masking. The Gaussian standard deviations were 0.2° visual angles on the inner side, and 1.2° visual angles on the outer side, resulting in faster tapering of the grating in the center, to stay clear of the fixation cross. The spatial frequency of the gratings was always 2 cycles per visual degree. The phase of the gratings was one of four values (0, *π*/2, *π*, 3*π*/2), which varied across trials, but was identical for S1 and S2 within each trial. The orientation of S1 was one of six values (15°, 45°, 75°, 105°, 135°, 165°). The orientation of S2 was determined by adding or subtracting the S1 orientation by the change size magnitude determined by the staircase procedure (see <u>Staircase procedure</u>).

Trial-by-trial co-variation in stimulus features (S1 orientation and grating phase), fixation duration, S1 duration (FP duration) and change direction (clockwise\counterclockwise) were determined in advance, to verify that they were maximally uncorrelated and vary independently (the correlation between each pair of features was |*r*_*pearson*_| < 0.014 in each of the FP conditions). In particular, this was crucial to ensure that FP duration was not predictable at the onset of S1 (other than the general knowledge about the FP distribution, collected over previous trials), and to ensure that the change direction in each trial was not predictable prior to the onset of S2. Additionally, we made sure that the trial type (change\no-change) was not predictable by any of the trial-level features (other than the duration of S1, which was always fixed to 1.85 sec for no-change trials; all other correlations |*r*_*pearson*_| < 0.007). Thus, the pairing of trial-level features was the same for all participants, but the order of trials was randomized per participant (see <u>Session procedure</u>).

### Staircase procedure

Task difficulty was determined by the magnitude of orientation difference between S1 and S2 (in absolute terms), which we refer to as the orientation tilt. The tilt was adjusted by an ongoing 1-up 3-down staircase procedure throughout the task, starting in the practice session.^63^ We used a logarithmic scale, with a starting tilt of 1, and linear step sizes of 1/3. In degree scale, this is equivalent to a starting tilt of roughly 2.72°, and a *multiplicative* step size of roughly 1.4. Having a 1-up 3-down procedure meant that following an incorrect response the tilt was multiplied by 1.4 (making the task easier) and following 3 consecutive correct responses the tilt was divided by 1.4 (making the task harder). The minimal and maximal tilts were set to 0.0013° and 39.1° respectively, and participants which arrived at these limits were excluded from further analysis (see <u>Participant exclusion</u>). The count towards 3 consecutive correct responses was restarted at breaks and end of practice but was not interrupted by no-change trials and missed response trials (see <u>Repeated trials</u>).

A 1-up 3-down staircase procedure is geared to identify the tilt magnitude resulting in 79.4% accuracy, which was indeed reached for the vast majority of participants (see <u>Results</u> and <u>Participant exclusion</u>). The staircase procedure was used to ensure a large enough dynamic range for detecting accuracy modulation by FP for all participants and avoid ceiling and floor effects (100% and 50% accuracy respectively). We kept the staircase procedure throughout the task to collect sufficient data while keeping the task (relatively) short, and to adjust for slower drifts in attention across the task.^64^ Tilt magnitude remained largely stable after the practice session (standard deviation of median tilt values across task blocks (mean ± SEM across participants in the main subject group): 0.21±0.02°, first block median tilt: 0.85±0.04°, last block median tilt: 0.71±0.04°).

### Session procedure

#### Instructions

Prior to performing the task participants were first provided with information about the task and gave their informed consent. Next, participants underwent a procedure to measure their screen size (see <u>Setup calibration</u>) and were provided with detailed instructions including examples of both trial types (change: 5 trials, 2 without responding and 3 with response, all with tilt magnitude of 7.39° (2 in logarithmic scale); no-change: 2 trials). Participants were requested to answer both accurately and quickly and were informed that their performance is measured on both accuracy and speed. Participants were told that the change magnitude will vary but were not informed regarding the relation of this to their performance. Additionally, they were informed that trials would be consecutive, and a short break will be provided every few minutes. We asked the participants to fixate on the central cross throughout the task (except for breaks), although we are not able to guarantee this was performed in the online setup. Participants were asked to prepare their index fingers over the ‘X’ and ‘N’ before starting the practice. Participants could review the instructions as many times as they wish.

#### Practice session

Following the instructions, participants performed a short practice session which included 16 trials (2 no-change). Tilt magnitude was initialized to 2.72°, after which it was changed adaptively in accordance with their performance (see <u>Staircase procedure</u>). S1 duration distribution was roughly uniform during the practice and the instructions examples. Following the practice, participants were provided with information about their performance (same as after each task block, see ‘Main task’ below). Participants were allowed to continue to the main task if they performed well in the practice, defined as achieving at least 70% accuracy rate and having no more than one out-of-time response of each type (too early or too late, see <u>Repeated trials</u>). Participants who did not perform well in the practice session were required to repeat the instructions and perform the practice again (until reaching the performance criterion, or until a maximum of 3 repetitions). Each time a participant failed to pass the practice the tilt magnitude was multiplied by 2.72 (3 step changes in the logarithmic scale) for the instruction’s examples and for the starting tilt of the next practice (e.g., after one repetition the tilt in examples was 20.09°, and the tilt at the start of the practice was 7.39°). From the main participant group, nearly all performed well in their first practice session (138/142, ∼97%). Most proceeded directly to the main task, while 2 chose to repeat the practice once more before proceeding. Before beginning the main task, participants were reminded to prepare their hands over the ‘X’ and ‘N’ keys, to maintain fixation during the task, and to keep a distance of approximately 60 cm from the screen (see <u>Setup calibration</u>).

#### Main task

The tilt magnitude at the start of the main task was identical to the tilt reached at the end of the practice session, to ensure that task difficulty was already adapted to each participant’s perceptual abilities (tilt median value for the first 10 change trials of the task (mean ± SEM across participants in the main group): 1.14° ± 0.06°, range: 0.51°-3.79°). The task consisted of 8 blocks, with 64-69 trials each (depending on the distribution condition), lasting roughly 3.5-4 min. Trials were assigned into blocks so that each trial-level feature was sampled with roughly equal proportions (see <u>Stimuli</u>). Trial order within each block was randomized per participant, with the only constraint being that no more than 3 no-change trials would be presented consecutively. Overall, sessions took approximately 45 minutes (mean ± SEM: 43 ± 0.5 min, range: 32 min to 1:17 hours), with the task part taking up approximately 80% of this time (mean ± SEM: 36 ± 0.3 min, range: 29-49 min).

#### Breaks

Between blocks participants were given short breaks, lasting up to 2 minutes (participants could elect to continue the task earlier by pressing the space bar). If the task was not actively resumed within 1:45 min a 15 second countdown clock would appear, and the task would resume automatically once it reached 0 sec. Participants varied in their mean break durations, from 2 seconds to 2 minutes (mean ± SEM across participants in the main group: 42.2 ± 2.4 sec). To facilitate engagement with the task, during breaks participants were provided with information about their performance in the previous block (accuracy, mean reaction time, and the number of trials with out-of-time responses; <u>Repeated trials</u>) and verbal feedback. Block feedback was positive if accuracy was above 70%, mean RT below 0.8 sec, and there were less than 3 out-of-time responses of each type (too early or too late). Otherwise, participants received feedback specific to the condition they did not satisfy.

### Repeated trials

During the main experiment, trials with responses that were administered too early (before the end of S2 for standard trials, or before the end of S1 for no-change trials) or too late (failed to respond within 2.5 sec of the onset of the question screen) were stopped and repeated at the end of the experiment in randomized order. In both cases feedback would be shown (‘Too soon, wait for the change’ or ‘Time is up, please respond faster’), and the task would pause for up to one minute or until the participant pressed the space bar (similar to the countdown clock displayed in the breaks, if the task was not actively resumed within 50 seconds a 10 second countdown clock would appear, and the task would resume once it reached 0 sec). If less than 12 trials had to be repeated these trials were added to the last block, otherwise, the last block would be followed by a short break and only then the repeated trials would begin. If during these trials, responses were again too early or too late that trial would be repeated again after all repeated trials of the previous round, until the participant responded in time. Most participants had just a few repeated trials (after participant exclusions, 83% of participants had 0-3 repeated trials; mean ± SEM across subjects: 1.8 ± 0.2, with roughly equal proportions for early presses and out of time responses).

### Setup calibration

As the experiment was administered online, we had no access to the precise setup participants were using. In the recruitment website, participants were instructed to use only a laptop or a desktop computer. However, the size of the screen is not available through the experiment website, therefore, participants performed a procedure to provide us with the information required to calibrate the stimulus presentation. The procedure, adapted from Morys-Carter’s screen scale code,^65^ takes advantage of the fact that credit cards have a consistent physical size across the world (85.6 mm x 53.98 mm). Participants are presented with a distorted image of a credit card and asked to hold up a physical credit card up to the screen and scale the image using the arrow keys until it is perfectly aligned with the real-life card. Thus, the final presentation scale of the image indicates how to present stimuli with a specific physical size, irrespective of the participant’s original setup. Screen sizes measured in this way were commensurate with participants complying with the requirements and using either a laptop or a desktop computer, except for one participant who was excluded from further analysis (see <u>Participant exclusion</u>; screen diagonal size mean ± SEM for the main subject group: 16.4 ± 0.4 inches, range 10.7-32.3 inch). To present stimuli at a consistent visual angle both the stimuli presentation size and the distance of the participant from the screen are required, therefore, we asked participants to sit 60 cm \ 23.5 inches away from the screen, and they were reminded of this before beginning the task.

## Analysis

### Data selection

Outlier trials were defined as trials with log-RT outside of the range of the mean ± 2 standard deviations of the entire log-RT distribution, including both correct and incorrect trials. We applied individualized thresholds, using for each participant their own log-transformed RT distribution (logarithm was applied to account for the skewness of RT distributions). These trials were excluded from both RT and accuracy analyses. Overall, this resulted in 21.1 ± 0.4 (mean ± SEM) trials excluded per participant (range: 4-35 trials). Considering only correct trials, we excluded 12.2 ± 0.3 (range: 2-23) trials per participant. Results without trial exclusion were similar to the main results.

### Behavioral quantification

To assess the influence of FP duration on behavior we examined the influence on perceptual accuracy (the percentage of correct trials) and correct trials RT (using the median RT as our main measure, due to the skewness of the RT distribution mentioned above). Single participant analyses focused on performance in the three shortest and three longest FPs (0.6-0.7 and 1.7-1.8 sec respectively; *Fig. 3a-b*). To get a robust single-subject measurement, we computed the accuracy and median correct RT for all three durations together (similar results were obtained by computing the accuracy and RT for each duration separately and then averaging). Computational modelling focused on the full behavioral time-courses (*Fig. 3c-d*) formed by computing the accuracy and median correct RT for each FP duration (for each participant separately, then averaged for each distribution condition).

### Statistical testing

Comparison of behavioral measurements within individuals across different time-points used paired t-tests per distribution, with Cohen’s d as the measure of effect size. This was done per distribution and corrected for multiple comparisons using Bonferroni correction. To compare behavioral measurements across distribution conditions we used one-way ANOVA, measuring effect size using partial eta squared. Post-hoc comparisons were performed using pairwise Tukey-HSD post-hoc tests.^66^ All tests were conducted using the pingouin package in Python.^67^

### Model fitting

To compare between the behavioral time-courses, constructed from the RT and accuracy of participants in the task (<u>Behavioral quantification</u>), and the computational models time-courses (<u>Computational Modelling</u>), we employed ordinary least squares (OLS) linear regression, implemented using the statsmodels package in Python.^68^ Linear regression was used to adjust for differences in units between the data and model time-courses. The time-courses of each data type (accuracy or RT) and each distribution condition were fit separately, to account for potential idiosyncrasies of specific participants (for how we combined results across distributions see Model evaluation).

For each empirical time-course and each model time-course, we identified the intercept 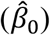 and scale 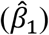 coefficients which most closely approximated the empirical time-course from the model time-course, by solving the following equation:

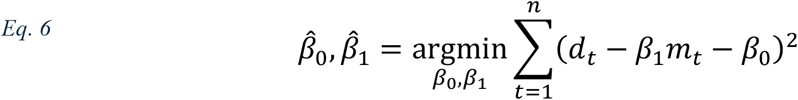

where *d*_*t*_ is the empirical behavioral data (accuracy\RT) at FP duration *t*, and *m*_*t*_ is the model prediction for the same FP. Both contained a total of *n* = 25 FP durations.

The model fits were then computed using the following equation:

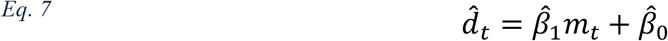

The computational models time-courses are intended to capture the dynamics of anticipation. Because higher anticipation (signified by higher model predictions), is expected to generate higher accuracy, but lower (faster) RT, the models are expected to have a *positive* relation to accuracy, and a *negative* relation to RT. To account for this, we restricted the sign of the scale coefficient 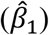 accordingly. If the solution to Eq. 6 did not satisfy the cognitive rational (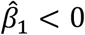 for accuracy data or 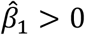 for RT data), we used a flat line as the model fit, as this is the best possible fit under the constraints (stemming from the convexity of Eq. 6). That is, we set 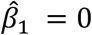 and solved the equation again, this time only for *β*_0_:

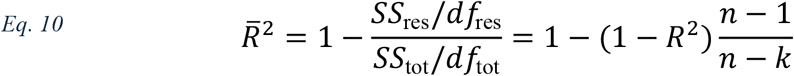

And the model fit was then given by:

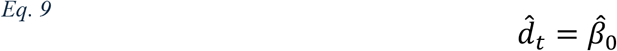

Additionally, in cases where the model prediction was constant for all time-points (e.g., PDF_UNI_ prediction), we used Eq. 8 and Eq. 9 directly, without attempting to fit a scale coefficient, since the model time-course in these cases was fully co-linear with the intercept predictor.

### Model evaluation

To evaluate the models we used adjusted coefficient of determination (adjusted R^2^, denoted by 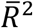), and Akaike Information Criterion (AIC).^26,27^ Both metrics quantify how well the computational models captured the variability in the behavioral data, while accounting for the number of predictors used in the regression.

Adjusted R^2^ was computed using the following equation:

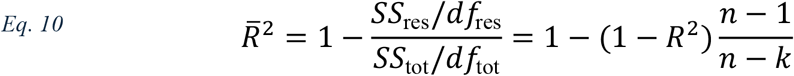

where *R*^2^ is the uncorrected coefficient of determination, quantifying the proportion of variance of the data that is explained by the computational model, *n* is the number of observations (in our case *n* = 25 FP durations in all models) and *k* is the number of parameters in the regression (*k* = 2 (intercept and slope) in all cases, including when 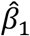 did not satisfy the cognitive constraints and the model was refit, except when model prediction was constant and then we used 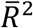 takes values between 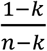 and 1 (closer to 1 indicating a better fit), and therefore for *k* = 2 the values of 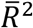 can be slightly negative.

AIC was computed using the following equation:

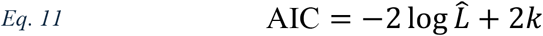

where 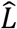 is the maximum likelihood of the regression, and *k* is the number of regression parameters (defined in the same way as for 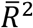). Importantly, ‘raw’ AIC values are not informative by themselves, as they are influenced by irrelevant factors such as the scale of the data (e.g., whether RT was measured in seconds or milliseconds). AIC provides a powerful theoretically motivated way to compare between models by computing the difference in AIC values between models (which is not influenced by irrelevant factors). Therefore, to compare between model A and model B we compute ΔAIC = AIC_*A*_ − AIC_*B*_ and examine the sign of this quantity. Models with *lower* AIC values are considered to be a better fit for the data (i.e., if ΔAIC < 0 model A is considered a better fit, and if ΔAIC > 0 model B is considered a better fit). The reason for this is that AIC is inversely related to the likelihood 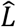, which quantifies how likely the data was to be observed given the model (and estimated scale and intercept coefficients). Therefore, models providing a better fit to the data have higher 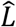 values, which leads to *lower AIC*. As is customary in the literature,^28^ we considered models to be substantially different in explaining the data only if 4 ≤ |ΔAIC|, while 2 ≤ |ΔAIC| ≤ 4 was considered a slight different in predictive ability, and models with |ΔAIC| ≤ 2 were not considered to be significantly different in explaining the data.

To combine results across distributions we used the sum of the AIC values obtained for each distribution separately (∑AIC). Summing AIC values in this way results in an identical AIC value to that which would be obtained by fitting all conditions together with condition-specific slope and intercept coefficients. This is because the likelihood 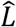 of a joint fit is equivalent to the product of 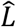 from each condition (since the data of each condition is independent) and the number of parameters *k* of a joint fit is equivalent to the sum of *k*s from each condition:

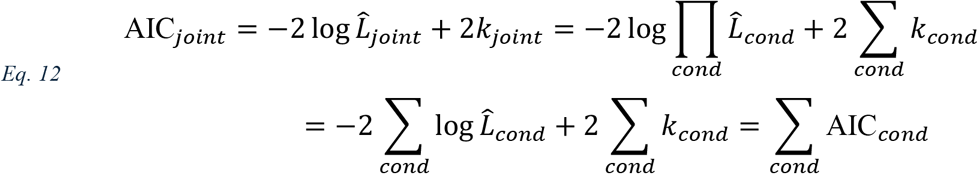

## Computational Modelling

### Hazard rate

The hazard rate of event *E* (HR_*E*_) at time *t* is defined as the probability that *E* will occur at *t*, given that it did not occur before *t*. Using Bayes rule we get:

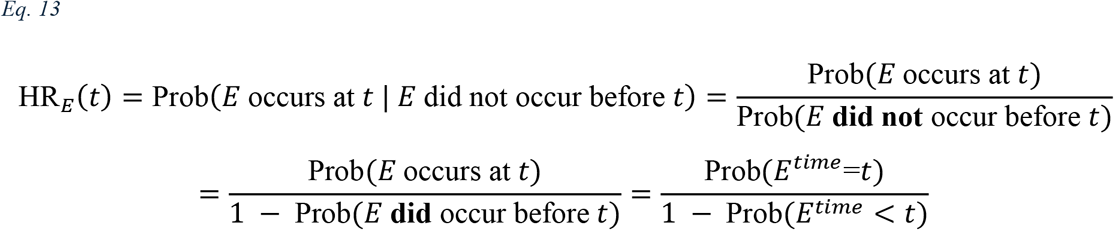

For continuous distributions, Prob(*E*^*time*^=*t*) is given by the PDF of *E*, which can be denoted by *f*(*t*), and Prob(*E*^*time*^ < *t*) is given by the cumulative distribution function (CDF), denoted by *F*(*t*). Therefore, we get:

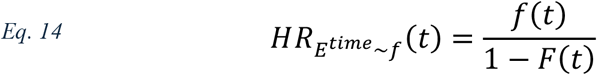

In our case the distributions were discretized by 50 ms intervals, therefore, in computing the model predictions we employed the HR computation for discrete random variables. Thus, we denote Prob(*E*^*time*^=*t*) by *p*(*t*), and 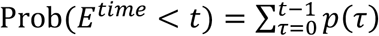 (the sum begins at 0, since time is non-negative), and together:

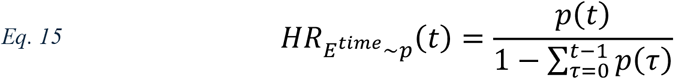

When the event always occurs, the denominator of this function approaches zero, which results in an infinite increase in the HR for long durations. In our case there were trials where the event never occurs (no-change trials, <u>Trial procedure</u>), and therefore the sum of the entire PDF is 0.875 (corresponding to 12.5% no-change rate), which limits the rise of the HR.

### ETemporal-estimation noise

People are not able to estimate elapsed time with perfect certainty,^24,25^ limiting their use of the task’s temporal structure. We modelled this by assuming that the subjective FP perceived in each trial is not the objective FP, but a noisy version of it. Formally, in trial *i*, with objective FP of *T*, the subjective FP 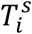 is given by:

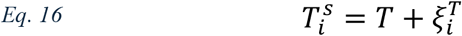

where 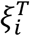 is normally distributed trial-specific noise:

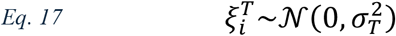

We assumed that the level of noise in the estimate may vary between different durations or between task contexts, and therefore, we denote the standard deviation of the noise distribution as *σ*_*T*_ and consider two possible factors for modulating this quantity (see <u>Noise modulation</u>). Additionally, we consider two separate influences of estimation noise on performance (see <u>Encoding and decoding noise</u>).

### Noise modulation

The level of noise in the estimate corresponds to the standard deviation of the noise distribution (*σ*_*T*_ in Eq. 17). Following ref. ^18–20^, we considered two hypotheses regarding the factor modulating this noise level: (A) Time-based modulation (modulation by the FP duration itself, *σ*_*T*_ ∝ *T*), and (B) Probability-based modulation (modulation by the frequency of the FP in the task, *σ*_*T*_ ∝ *f*(*T*)).

#### Time-based modulation

A large body of work has shown that in timing related tasks such as temporal reproduction the ratio between the duration being reproduced, and the standard deviation of the reproduction is roughly constant. This constant is known as the Weber fraction, or the ‘scalar property’ of time.^24,25,29^ In other words, the noise in temporal estimation is proportional to the duration of the estimate. Therefore, in our central investigation we modelled *σ*_*T*_ as a linear function of the FP duration *T*:

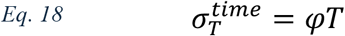

where *φ* denotes the Weber fraction. We set *φ* = 0.21 in all of our models, in accordance with prior work.^18–20^

#### Probability-based modulation

An alternative hypothesis, promoted recently by Grabenhorst and colleagues,^18–20^ is that estimation noise depends not on the FP duration itself, but on the probability of each FP duration (given by the PDF, *f*(*T*)), such that FPs with high probability are estimated with *less* noise. Thus, *σ*_*T*_ are assumed to be a linear function of −*f*(*T*), scaled to match the range of noise levels obtained by time-based modulation. For the EXP and FLIP-EXP conditions, the following equation is used:

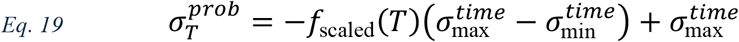

with 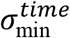 and 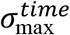 denoting the minimum and maximum noise level values under time-based modulation (0.6 ⋅ *φ* and 1.8 ⋅ *φ* respectively), and *f*_scaled_(*T*) denoting the PDF scaled between 0 and 1:

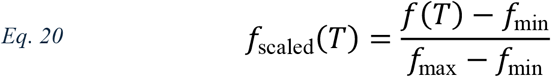

For the UNI condition (constant PDF) 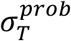 was set to the mean of 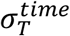 (1.2 ⋅ *φ*) for all FPs.

### Encoding and decoding noise

We hypothesized that temporal-estimation noise can have two distinct influences on the task responses, stemming from the two downstream uses of the subjective FP estimate in each trial. These were added to our model as two distinct computational stages (*Fig. 1b*):

A. ‘Encoding’ – using the current FP estimate to adjust the internal representation of the task temporal structure (that is, to update the subjective PDF representation,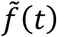). This impacts the anticipation which is used to guide behavior in future trials. If encoding is noise-free, the subjective FP estimate is identical to the objective FP (denoted by *T*), and if encoding is noisy, the estimate is equal to 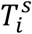, given by Eq. 16.
B. ‘Decoding’ – using the FP estimate to infer or calibrate the current anticipation level, which modulates response in the current trial. As anticipation is time-dependent, this can be formalized as evaluating a temporal function (denoted here as 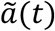). Similarly to the encoding case, if decoding is noise-free the subjective FP is identical to the objective one, so that the anticipation modulating the response in the current trial is 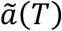, while for noisy decoding the anticipation is given by 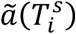.

In this formulation, ‘Encoding’ and ‘Decoding’ are both trial-level processes. To form the model-predicted time-courses, we examine their influence at the task-level, by computing their expected influence (in the probability sense) across trials. Moving to the task-level is also crucial to discern the influence of the noise on each stage, as the noise on each trial is stochastic, but the distribution-level effects follow a predictable structure. Framed this way, task-level encoding can be conceptualized as transforming the objective PDF *f*(*t*) to the subjectively encoded PDF 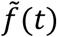, and task-level decoding as a transformation from the internal subjective anticipation 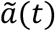 to the information *a*(*t*) which was used in practice to guide behavior. Importantly, we do not mean to infer that each of these functions is represented explicitly in the brain. While this may be the case for the subjective PDF 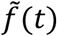 and the subjective anticipation 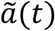 (although other alternatives also exist, see <u>Discussion</u>), the decoded anticipation *a*(*t*) is a mathematical construct generated to enable comparison of the model predictions to the empirical data.

Formally, the task-level ‘Encoding’ and ‘Decoding’ for each duration *t* are defined as follows:

A. The encoded PDF 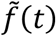 is the expected probability of estimating subjectively a trial’s FP duration as *t*, i.e., the probability that any trial (with any *objective* FP, denoted by *T*^*o*^, distributed according to the objective PDF *f*) was estimated *subjectively* as having a duration of *t* (*T*^*s*^ = *t*). From the law of total probability:

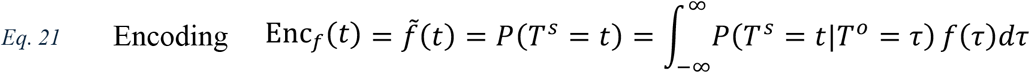
B. The decoded anticipation *a*(*t*) is the expected subjective anticipation 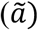 for a trial with objective FP duration *t*. That is, *a*(*t*) is given by expectation of 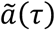, the internal subjective anticipation, across all trials estimated *subjectively* as *τ* (*T*^*s*^ = *τ*), given that the *objective* FP was *t* (*T*^*o*^ = *t*):

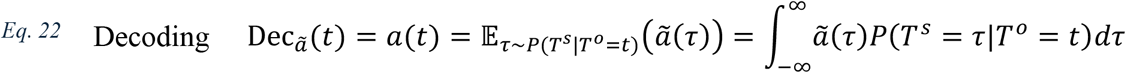

For noise-free encoding and decoding, *P*(*T*^*s*^ = *t*|*T*^*o*^ = *τ*) and *P*(*T*^*s*^ = *τ*|*T*^*o*^ = *t*) are 1 for *t* = *τ* and 0 otherwise. Therefore, in this case, both are equal to identity transformations:

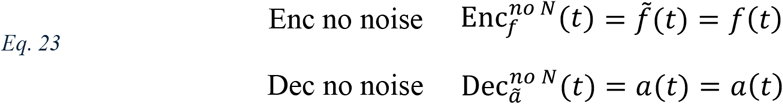

To examine the influence of noise on both stages, note that from Eq. 16 and Eq. 17 it follows that the probability of a trial with *objective* FP duration of *T*^*o*^ = *o*, being perceived *subjectively* as having an FP duration of *T*^*s*^ = *s*, is given by:

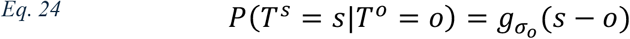

where *g*_*σ*_(*x*) denotes a Gaussian distribution with zero mean and standard deviation of *σ*:

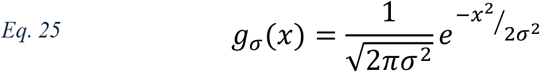

Substituting this in Eq. 21 and Eq. 22, we get:

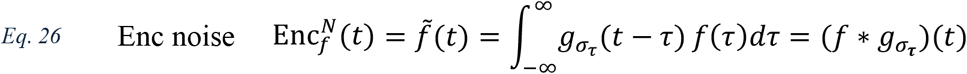

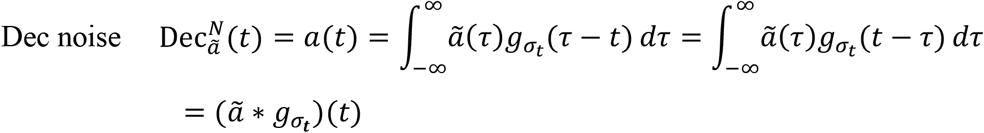

where ∗ denotes convolution. Therefore, the influence of both encoding and decoding noise is akin to convolving a temporally resolved input function with a Gaussian kernel, with two important differences shown in Eq. 26: (1) The temporally resolved input function – encoding acts on the objective PDF *f*, which decoding acts on the internal subjective anticipation 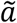, and (2) Which duration underlies *σ* the level of noise – in encoding each duration in the objective PDF is impacted by its own noise level, therefore the encoded PDF combines many different noise sources (we received that *σ*_*enc*_ = *σ*_*τ*_, where *τ* is the duration being integrated in the convolution). In contrast, for decoding the noise level is determined only by the decoded duration, therefore it is identical for each subjective duration being integrated (we received that *σ*_*dec*_ = *σ*_*t*_, where *t* is the output of the convolution).

All models were evaluated at 50 ms intervals, keeping with the 50 ms discretization of the FP distributions. In the models including noise at the encoding or decoding stages, we approximated the integral in the convolution (Eq. 26) using the trapezoid rule (implemented in scipy)^69^ with 50 ms discretization for *dτ* and zero-padding on both sizes of the input function to adjust for edge effects.

### Nonlinear transformations

We considered three nonlinear transformations on the internal representation: (1) Logarithm (2) Reciprocal (3) Exponential. All three were incorporated into our modelling framework following the PDF or HR calculation (depending on the model) and before the decoding stage (*Fig. 1b*). As both the PDF and the HR are probabilities (corresponding to the a-priori probability of each FP, or the conditional probability, respectively), the input to the transformations was always in probability form and will be denoted by *p*.

#### Logarithmic

models predict that anticipation to an FP of probability *p* corresponds to log(*p*). We tested this because it features broadly in psychophysical explanations^33,34^ and neural coding schemes,^31,32,35^ and because of the relation of this function to Shannon’s information theory,^36^ where 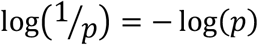 quantifies the information content of an event of probability *p* (also known as ‘Shannon surprisal’). Therefore, both the PDF-log and the HR-log models predict that the anticipation to a specific FP is not related to the probability (or conditional probability) of this FP, but inversely to how surprising it is, so that more surprising FPs are anticipated less.

#### Reciprocal

models predict that anticipation to an FP of probability *p* corresponds to the negative of the reciprocal, − 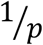. The reciprocal model was tested due to recent suggestions in the literature that the reciprocal of the PDF underlies temporal anticipation.^18–20^ This model suggests that anticipation to a specific FP is (inversely) related to the average number of trials that occur before a trial of duration FP is encountered. Thus, FPs with lower probability (requiring a longer wait), are anticipated less.

#### Exponential

models predict that anticipation to an FP of probability *p* corresponds to exp(*p*). These were tested for theoretical completeness. Since both the logarithmic and the reciprocal transformations are concave functions (within the relevant range of probabilities), with greater emphasis on lower probabilities, we added a convex operation to the modelling framework. The exponential was ideally suited for this as it is mathematically complementary to the logarithm.

## Code

The task was implemented in JavaScript using PsychoJS, created using Psychopy builder with code components (version 2023.2.3),^70^ and hosted online using the Pavlovia platform (https://pavlovia.org/). Analysis was performed using custom code in Python (version 3.11), relying heavily on pandas^71^ and numpy^72^ functionality, as well as matplotlib^73^ and seaborn^74^ for visualization, and other libraries where noted. Analysis and experiment code will be made available upon review.

## Acknowledgements

We are grateful to N. Ofir for insightful comments about the manuscript. We also thank members of the Human Cognitive Neuroscience Laboratory (HCNL) and members of the Brain, Attention & Time Lab for support throughout the study. G.V. is grateful to S. Ravfogel for his support throughout the study and especially in writing the manuscript, and H. Vishne for assistance with figures. In work on this study G.V. was supported by the Azrieli Foundation graduate fellowship, the Rothschild Postdoctoral Fellowship and the Zuckerman STEM leadership program. L.Y.D. is supported by the Jack H. Skirball research fund and by Israel Science Foundation (ISF) grant 3504/20. The Brain Attention & Time Lab (PI: A.N.L.) is supported by the James McDonnell Scholar Award in Understanding Human Cognition and ISF grant 958/16. This project has received funding from the European Research Council (ERC) under the European Union’s Horizon 2020 research and innovation programme (grant agreement no. 852387).

## Author contributions

Conceptualization and methodology: G.V., L.Y.D. and A.N.L. Investigation, data curation, formal analysis and software: G.V. Funding acquisition and supervision: L.Y.D. and A.N.L. Writing – original draft and original visualization: G.V. Writing – review and editing and visualization input: G.V., L.Y.D. and A.N.L.

